# An Enantioselective Chemical Probe for Chikungunya nsP2 Helicase with Antialphaviral Activity

**DOI:** 10.1101/2025.04.24.650479

**Authors:** Bose Muthu Ramalingam, Hans J. Oh, John D. Sears, Chun-Hsing Chen, Anand Vala, Shubin Liu, Kacey M. Talbot, Mohammed Anwar Hossain, Peter J. Brown, Scott Houliston, Julia Garcia Perez, Fengling Li, Meareg G. Amare, Peter Halfmann, Jessica L. Smith, Alec J. Hirsch, Cheryl H. Arrowsmith, Levon Halabelian, Ava Vargason, Rafael M. Couñago, Jamie J. Arnold, Craig E. Cameron, Nathaniel J. Moorman, Mark T. Heise, Timothy M. Willson

**Affiliations:** Structural Genomics Consortium, UNC Eshelman School of Pharmacy, University of North Carolina at Chapel Hill, Chapel Hill, NC 27599, USA; READDI AViDD Center, University of North Carolina at Chapel Hill, Chapel Hill, NC 27599, USA; Department of Microbiology and Immunology, University of North Carolina at Chapel Hill, Chapel Hill, NC, 27599, USA; X-Ray Core Laboratory, Department of Chemistry, University of North Carolina at Chapel Hill, Chapel Hill, NC 27514, USA; Piramal Discovery Solutions, Pharmaceutical Special Economic Zone, Sarkhej, Bavla Highway, Ahmedabad, Gujarat 382213, India; Research Computing Center, University of North Carolina, Chapel Hill, NC 27599, USA; Structural Genomics Consortium, University of Toronto, Toronto, Ontario, M5G 1L7, Canada; Princess Margaret Cancer Centre, University Health Network, Toronto, Ontario, M5G 1L7, Canada; Department of Pathobiological Sciences, School of Veterinary Medicine, University of Wisconsin, Madison, WI 53706, USA; Vaccine and Gene Therapy Institute, Oregon Health & Science University, Beaverton, OR 97006, USA; Department of Medical Biophysics, University of Toronto, Toronto, ON, Canada; Department of Pharmacology and Toxicology, University of Toronto, Toronto, ON, Canada; Eshelman Innovation, UNC Eshelman School of Pharmacy, University of North Carolina at Chapel Hill, Chapel Hill, NC, 27599, USA; Center of Medicinal Chemistry, Center for Molecular Biology and Genetic Engineering, University of Campinas, 13083-886-Campinas, SP, Brazil; Department of Genetics, University of North Carolina at Chapel Hill, Chapel Hill, NC, 27599, USA

**Keywords:** helicase, alphavirus, inhibitor, enantioselective, chemical probe, allosteric

## Abstract

Chikungunya virus (CHIKV) replication relies on the multifunctional nsP2 protein, making it an attractive target for antiviral drug discovery. Here, we report the resolution of oxaspiropiperidine **1**, a first-in-class inhibitor of the CHIKV nsP2 RNA helicase (nsP2hel), into its constitutive enantiomers and characterization of their antiviral activity. The enantiomer (*R*)-**1** exhibited potent inhibition of viral replication, nsP2hel ATPase activity, and dsRNA unwinding, while the (*S*)-**1** enantiomer was >100-fold less active. The (*R*)-**1** enantiomer also demonstrated high selectivity for CHIKV over other RNA viruses and for nsP2hel over other RNA helicases. Direct binding of (*R*)-**1** to nsP2hel protein was confirmed by ^19^F NMR. Biophysical and structural studies revealed conformational polymorphism in the spirocyclic scaffold of (*R*)-**1**, suggesting a potential role of thermal mobility of the ligand in allosteric inhibition of nsP2hel. Collectively, these findings designate (*R*)-**1** (RA-NSP2-1) as a high-quality chemical probe and (*S*)-**1** (RA-NSP2-1N) as a negative control for probing the biology of alphavirus RNA helicases.

---

Alphaviruses are mosquito-borne RNA viruses that pose a significant public health risk both within the United States and globally.^1^ Alphaviruses such as Chikungunya virus (CHIKV), Ross River virus, and O’nyong-nyong virus have caused large scale epidemics of debilitating acute and chronic arthralgia. Likewise, Venezuelan (VEEV), Western, and Eastern (EEEV) Equine Encephalitis viruses trigger neurologic disease with high rates of morbidity and mortality^2^ with a recent outbreak of EEEV in New England resulting in issuance of multiple public health alerts.^3^ The threat posed by these viruses is likely to increase as global warming expands the footprint of their mosquito vectors. Moreover, viruses such as VEEV and EEEV are also considered potential bioterror threats due to their capacity for weaponization.^4^ Despite the public health danger posed by these viruses, there are currently no FDA-approved drugs for any alphavirus-caused disease. Broadly acting antiviral therapeutics are badly needed for protection against both existing and future alphavirus threats.

RNA helicases are a diverse class of ATP-dependent motor proteins that play essential roles in nearly every aspect of RNA metabolism, including transcription, splicing, translation, RNA transport, and decay.^5, 6^ By unwinding RNA duplexes or remodeling ribonucleoprotein complexes, helicases modulate RNA structures critical for cellular and viral gene expression. The superfamily 1B (SF1B) helicases, characterized by conserved RecA-like domains, operate through nucleotide-induced conformational changes to translocate along RNA substrates in a 5’-3’ directional manner.^7^ Viral RNA helicases of have emerged as key targets for therapeutic intervention due to their indispensable roles in genome replication and virulence. Alphaviruses encode an SF1B RNA helicase within the N-terminal domain of non-structural protein 2 (nsP2hel) that is required for synthesis of viral RNA. In addition to the helicase domain, nsP2 also contains a cysteine protease domain which is connected to a C-terminal methyltransferase-like (MTL) domain.^8^ The alphavirus nsP2hel is a dynamic nonprocessive motor enzyme that detaches and re-binds as it unwinds the RNA. Recent studies suggest that the C-terminal MTL domain may stabilize the RNA-nsP2 interaction and is therefore required for efficient helicase unwinding activity.^9^ Given its essential role in alphavirus replication, nsP2hel is a promising and target for antiviral drug discovery. However, despite increasing structural knowledge from both X-ray crystallography^10^ and cryoEM studies,^11^ the development of selective small molecule inhibitors for this dynamic and conformationally flexible enzyme has proven challenging and few chemical tools exist to probe nsP2hel function or to enable mechanistic study of its activity.

The oxaspiropiperidine **1** (Figure 1) was recently reported as a first-in-class nsP2hel inhibitor with broad spectrum antialphaviral activity.^12^ Oxaspiropiperidine **1** was shown to be a direct-acting inhibitor of CHIKV nsP2 by the mapping of viral resistance mutants to the RecA1 domain of the helicase and with a difluoro analog demonstrating binding to nsP2hel protein by ^19^F NMR.^12^ Inhibition of nsP2hel was non-competitive with either the ssRNA substrate or ATP cofactor indicating that **1** was likely to be a Type 1 allosteric inhibitor.^13^ Notably, **1** contains a chiral center at C-4 but was tested initially as a racemic mixture of enantiomers. In this report, we describe the resolution of **1** at C-4, the dramatic difference in enzyme inhibition and antiviral activity between its two enantiomers, and the assignment of absolute configuration of the active enantiomer as (*R*)-**1**. We also show evidence of thermal mobility in the piperidine ring that may be an important property of this RNA helicase inhibitor. The active enantiomer (*R*)-**1** meets the criteria of a high-quality chemical probe (designated RA-NSP2-1) for CHIKV nsP2hel, while the inactive (*S*)-**1** enantiomer (designated RA-NSP2-1N) functions as a negative control analog.

**Figure 1.**
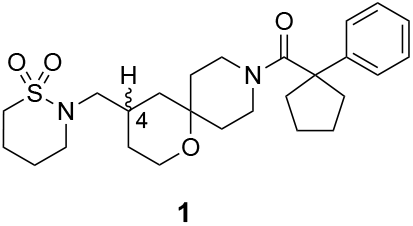
Racemic oxaspiropiperidine nsP2hel inhibitor **1**. The chiral center at C-4 is indicated.

## Results and Discussion

### Resolution and Enantioselectivity of Oxaspiropiperidine 1

The racemic oxaspiropiperidine **1** underwent separation by supercritical fluid chromatography (SFC) on an amylose-based Chiralpak AD-H column using CO_2_ as the primary mobile phase to yield the individual enantiomers. The enantiomers were initially designated (+)-**1** and (-)-**1** using their optical rotations (Table 1). Analytical HPLC using a polysaccharide Chiralpak IC column monitoring at 220 nm indicated that each enantiomer was isolated at >98% enantiomeric excess (ee). Testing of the individual enantiomers obtained by chiral SFC in the CHIKV-nLuc viral replication assay^12^ showed that antiviral activity was retained in (+)-**1** (Table 1). The (+)-**1** enantiomer was slightly more potent than racemic (±)-**1** and ∼300-fold more active than the (-)-**1** enantiomer. Thus, the antiviral activity of **1** showed remarkable enantioselectivity, with activity residing dominantly in the (+)-**1** enantiomer.

**Table 1.**
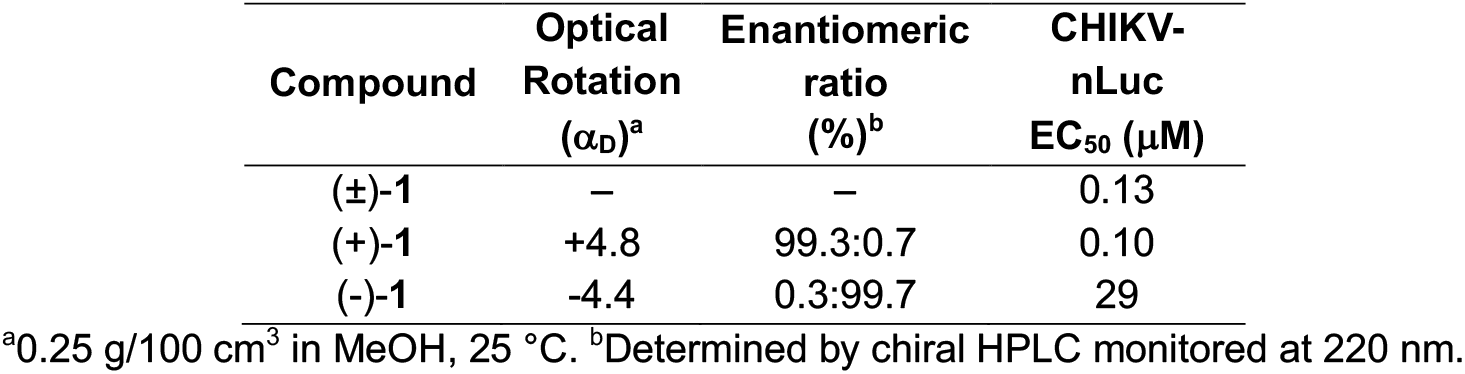
Antialphaviral Activity of 1 and its enantiomers.

### Assignment of absolute configuration

Kinetic resolution of a synthetic intermediate **2** (Figure 2) was used to produce larger quantities of the individual enantiomers of **1** for further biological characterization and to enable assignment of absolute configuration at C-4. Several chiral carboxylic acids were screened to obtain salts of racemic intermediate **2** containing a primary aminomethylene at C-4 of the oxaspiropiperidine core (Figure 2). Diastereomeric salt formation was successful using di-*p*-toluoyl-D-tartaric acid (D-DTTA) in EtOH at 90 °C. Upon cooling to room temperature, a crystalline salt composed of **2** with D-DTTA was formed in a 2:1 ratio as determined by ^1^H NMR (Figure S1). The crystalline salt was collected by filtration and washed with EtOH in 40% total yield (80% yield of a single enantiomer). Chiral HPLC analysis of the initial batch of (**2**)_2_•D-DTTA salt showed predominantly a single isomer with >95:5 enantiomeric ratio. Recrystallization of the initial (**2**)_2_•D-DTTA salt from EtOH on a gram scale further enriched the dominant enantiomer. Confirmation of the enantiomeric ratio (98.3:1.7, > 96% ee) was obtained by chiral HPLC analysis following conversion of **2** to a *p*-methoxybenzoyl derivative **3** that contained a strong UV chromophore at 254 nm (Figure 2).

**Figure 2.**
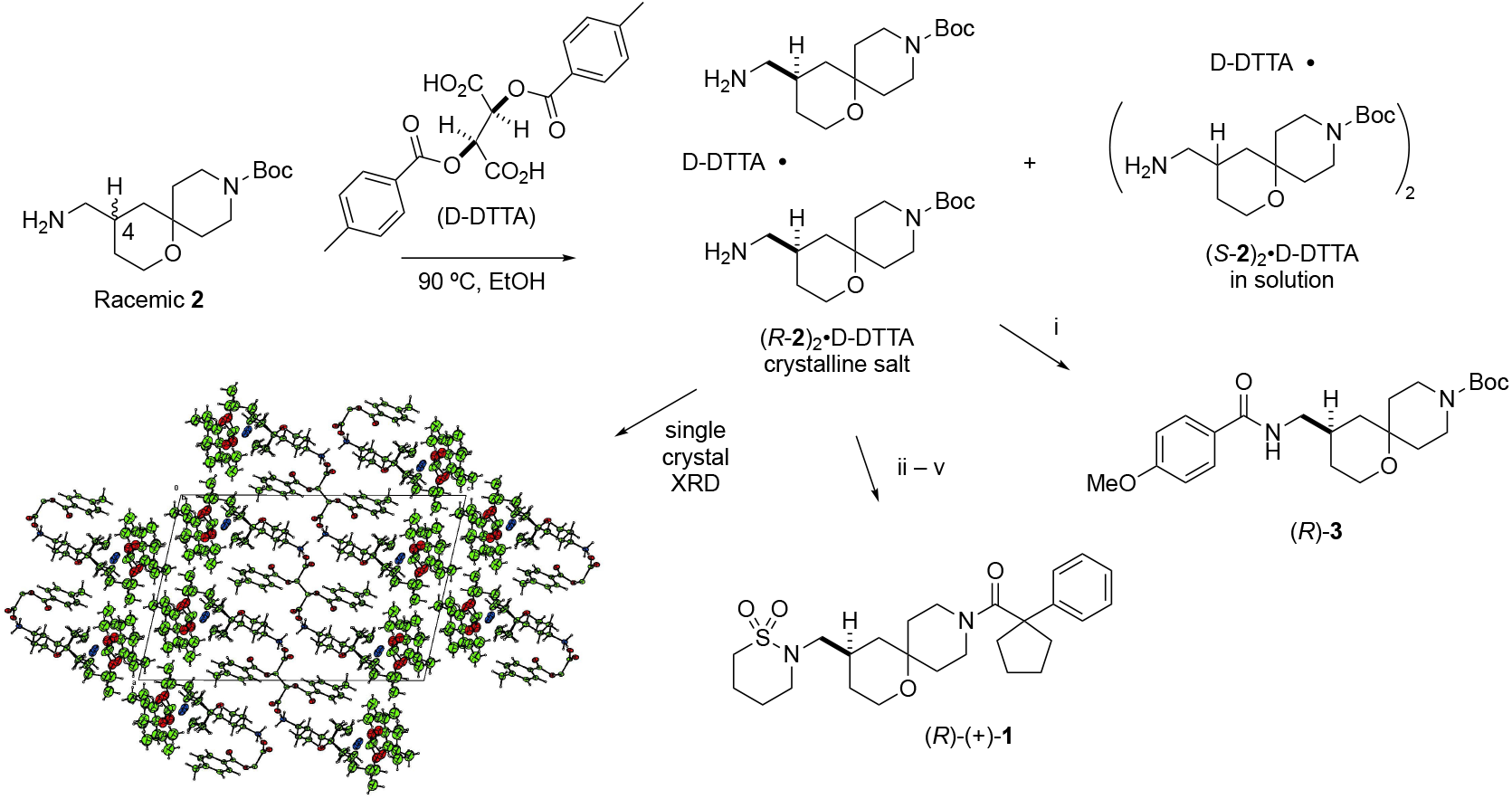
Assignment of absolute configuration at C-4 of the active enantiomer. (i) 1M KOH solution, rt; then 4-methoxybenzoyl chloride, Et_3_N, DCM; (ii) 1M KOH solution, rt; then 4-chloro-1-butylsulfonyl chloride, Et_3_N, DMF; (iii) DBU, ACN, 70 ºC; (iv) HCl in dioxane; (v) 1-phenylcyclopentane-1-carboxylic acid, EDC, HOBT, DIPEA, DMF.

To determine the absolute configuration at C-4 of the active enantiomer single crystal X-ray diffraction (SC-XRD) analysis was performed on a single crystal of (**2**)_2_•D-DTTA grown in methanol. Although the X-ray structure had a complex unit cell due to multiple conformations of the Boc group (Figure 2 and Tables S1–S6), the tetrahydropyran ring was well resolved and revealed that the (**2**)_2_•D-DTTA salt had the *R*-configuration at C-4 (Figure 2 and Figure S2). A four-step sequence was used to convert (*R*-**2**)_2_•D-DTTA into (*R*)-**1**, which demonstrated > 99% ee by chiral HPLC analysis at 230 nm. Measurement of the optical rotation of (*R*)-**1** showed it to be the dextrorotatory (+)-isomer. Thus, the absolute configuration of the active enantiomer with antiviral activity was determined as (*R*)-(+)-**1**. Using a parallel sequence of reactions, kinetic resolution of intermediate **2** using di-*p*-toluoyl-L-tartaric acid (L-DTTA) yielded (*S*-**2**)_2_•L-DTTA after recrystallization. Following the four-step sequence from (*S*-**2**)_2_•L-DTTA, the inactive (*S*)-(-)-**1** enantiomer was obtained with > 99% ee as determined by chiral HPLC analysis at 220 nm.

### Conformational Polymorphism of (R)-1

Notably, neither racemic **1** or (*R*)-**1** were crystalline. Even on a gram scale (*R*)-**1** was isolated a stable non-crystalline amorphous powder (Figure 3A). Compounds that exist in multiple conformations often do not form a stable crystal lattice and are resistant to crystallization, a property described as conformational polymorphism.^14^ To date no successful strategy has been found to induce (*R*)-**1** to crystallize even after multiple attempts over several months. The NMR spectra of **1** and several close analogs previously indicated that the oxaspiropiperidine core often did not exist in a single conformation in solution.^12^ Likewise, the solution ^1^H and ^13^C-NMR spectra of (*R*)-**1** showed evidence of conformational polymorphism with broad signals observed for the piperidine ring at room temperature (Figures 3B and 3C). Assignment of the NMR spectra was made by reference to an analog that was studied by variable temperature in multiple solvents.^12^ In addition, as noted above, refinement of the SC-XRD structure of (*R*)-**2** (Figure 3D) which had a single crystal morphology and packing showed a two-part disorder in the piperidine ring with each component exhibiting a further two-part disorder of the Boc group (Figure 3E). While the overall solution of the X-ray structure did not change, thermal motion in the oxaspiropiperidine core resulted in disorder in the electron density for the piperidine ring and Boc group (Figure S2). Together, these data indicate that **1** and its enantiomers are likely to exist as conformational isomers in both the solid state and in solution.

**Figure 3.**
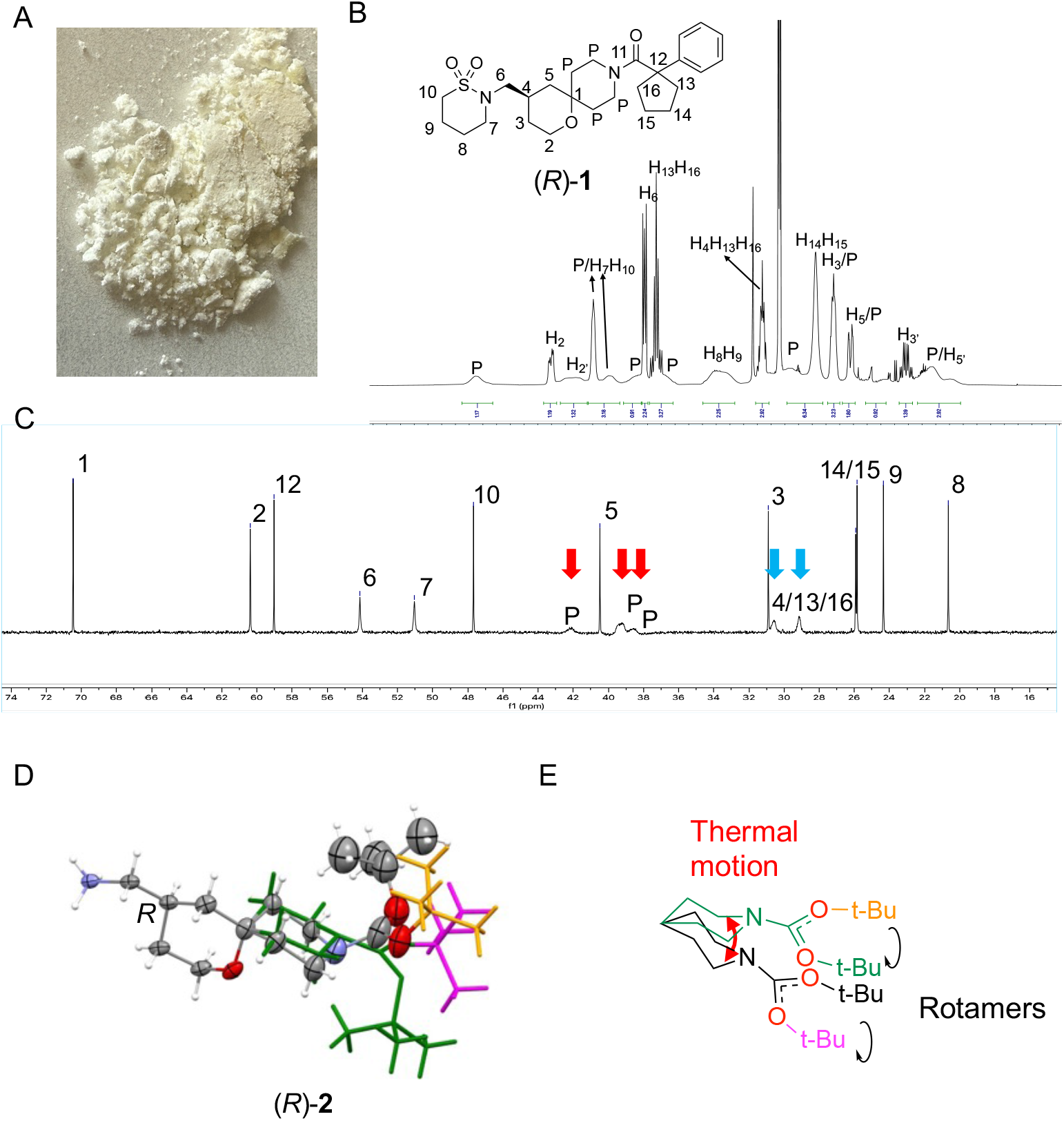
Conformational polymorphism and thermal motion of the oxaspiropiperidine ring. (A) (*R*)-**1** was isolated as a stable non-crystalline powder. (B) ^1^H NMR spectrum of (*R*)-**1** in CD_3_CN at 25 ºC showing line broadening of the piperidine protons (P). (C) ^13^C NMR spectrum of (*R*)-**1** in pyridine-*d*_5_ at 25 ºC showing line broadening of the piperidine carbons (red arrows) and cyclopentane carbons (blue arrows). (D) Refined X-ray structure of (*R*)-**2** showing disorder in the piperidine ring and Boc group. (E) Depiction of the of thermal motion in the piperidine ring and the amide rotamers seen in the SC-XRD structure of (*R*)-**2**.

To further understand the energetic barrier to motion in the oxaspiropiperidine core, *ab initio* DFT calculations were performed on the (*R*-**2**)_2_•D-DTTA salt to determine the interaction energy between the molecules following full structure optimization. Starting from the four limiting conformations observed in the X-ray coordinates (Tables S2–S6) as the initial geometries (Figure S3), the minimized structures converged on two amide rotamers of the Boc group that differed by 0.9 kcalmol^-1^ (Figure S4) with only a single minimized conformation for the piperidine ring, possibly due to a low barrier to motion in the ring. These gas phase calculations combined with the experimental solution NMR and SC-XRD structure indicate that thermal motion in the piperidine ring is a property of the spirocyclic core and is linked to the conformational polymorphism of the amide rotamers.

### Enantioselectivity in nsP2 Helicase Inhibition

To confirm that enantioselective antiviral activity (Table 1) was due to differences in the inhibition of nsP2 helicase by (*R*)-**1** and (*S*)-**1**, biochemical ATPase and unwindase assays were performed. Both biochemical assays utilized CHIKV nsP2 protein containing the helicase, protease, and methyl transferase-like domains, since it was known that the full-length protein was required for efficient RNA strand separation activity.^9^ ATPase activity was determined using a coupled ADPglo assay that monitored ADP production catalyzed by nsP2.^12^ In this assay, (*R*)-**1** inhibited ATPase activity with an IC_50_ = 0.84 μM with (*S*)-**1** showing >100-fold weaker potency for enzyme inhibition (Figure 4A). A FRET-based strand separation assay using an RNA duplex was used to measure unwindase activity of nsP2. (*R*)-**1** showed a IC_50_ = 130 nM for RNA unwindase inhibition and was 200-fold more potent than (*S*)-**1** (IC_50_ = 26 μM, Figure 4B). Thus, the nsP2-dependent ATPase and unwindase inhibition of (*R*)-(+)-**1** and (*S*)-(-)-**1** matched their antiviral activity in potency and enantioselectivity providing strong support for the cellular effects being mediated through disruption of the helicase enzyme.

**Figure 4.**
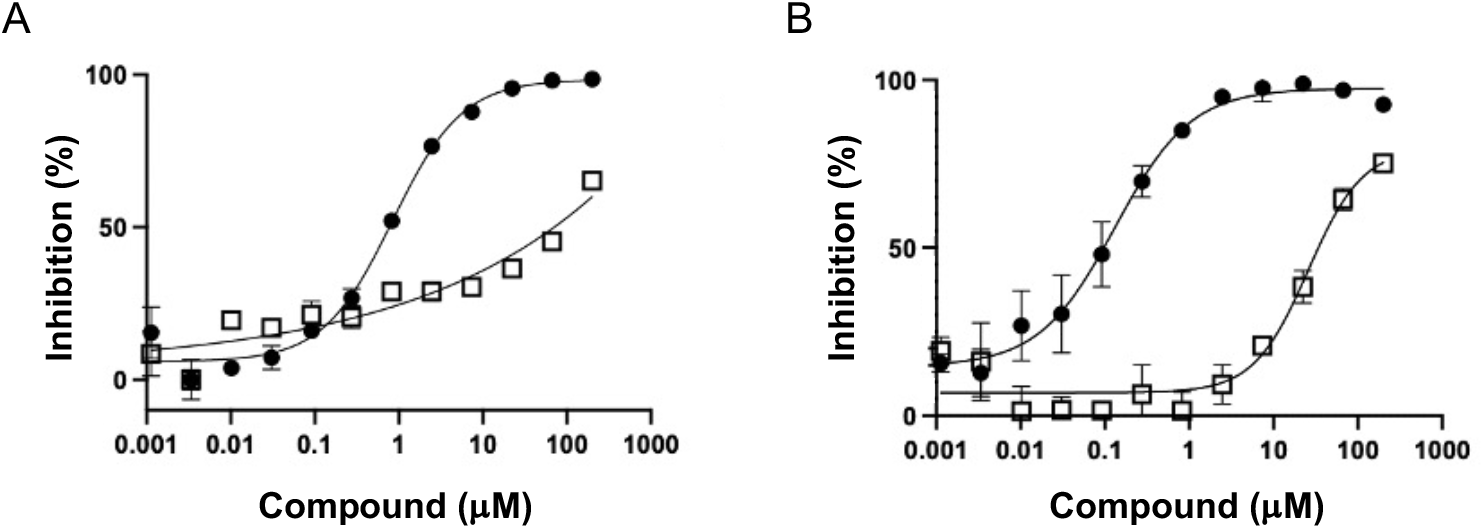
Inhibition of nsP2 helicase by (*R*)-**1** (closed circles) and (S)-**1** (open squares). (A) ATPase assay. (B) RNA unwinding assay. Error bars represent the average of triplicate determinations.

### Target Engagement

To assess direct binding to the CHIKV nsP2hel, the (*R*)- and (*S*)-enantiomers of difluorinated oxaspiropiperidine **4** were synthesized (Figure 5A). (*R*)-**4** demonstrated >100-fold greater potency than (*S*)-**4** in the CHIKV antiviral, ATPase, and RNA helicase unwinding assays (Figure S5) matching the enantioselectivity in biological activity that was seen with **2**. To demonstrate binding the ^19^F NMR of (*R*)-**4** titrated with nsP2hel protein was measured. The spectra revealed a dose-dependent decrease in ^19^F peak area with increasing nsP2hel protein, indicative of a binding interaction between the inhibitor and the protein (Figure 5B). In contrast, (*S*)-**4** showed no decrease in ^19^F peak area in the presence of nsP2hel protein. Importantly, the enantioselective binding of **4** to nsP2hel aligned with the enantioselective antiviral and biochemical activity.

**Figure 5.**
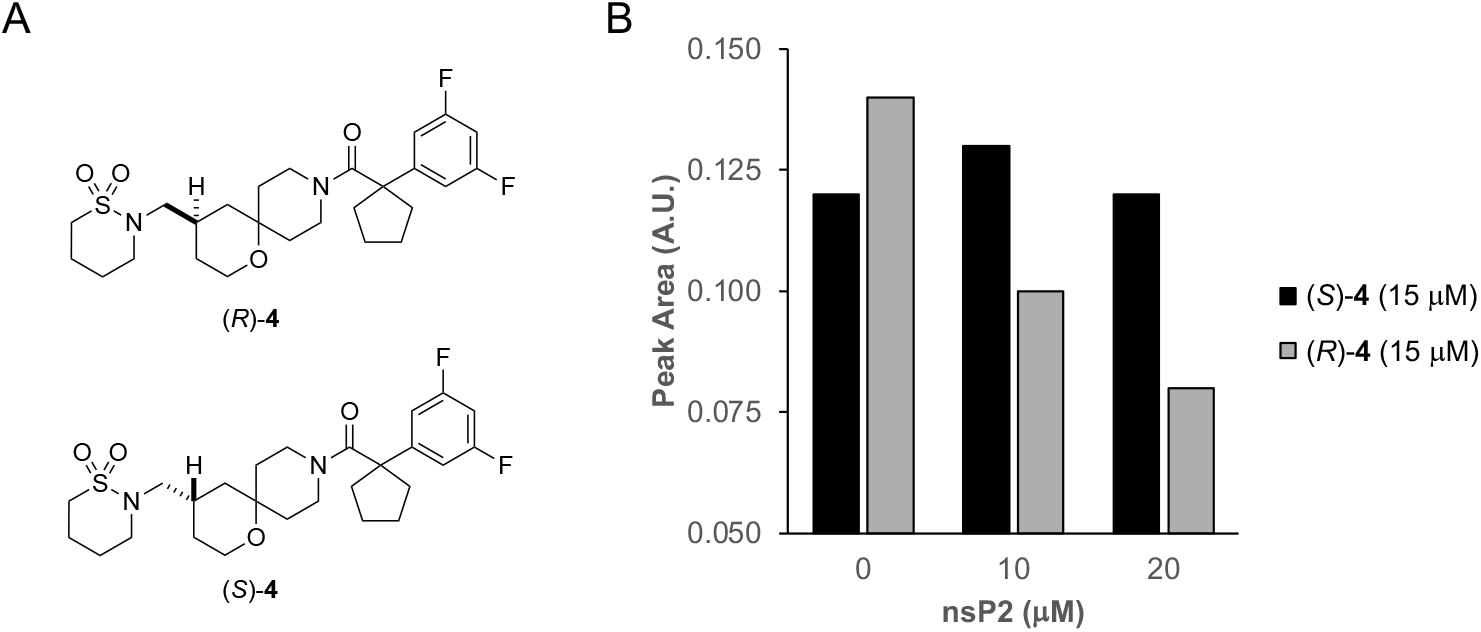
Target engagement by ^19^F NMR. (A) Difluorinated analogs (*R*)-**4** and (*S*)-**4**. (B) ^19^F NMR peak areas of (*R*)-**4** and (*S*)-**4** in the absence or presence of nsP2hel protein (aa 1–464). Relative concentrations of the ligand and protein are indicated.

### Target Selectivity

To assess the target selectivity of **1**, both enantiomers were tested for activity on five RNA viruses using luciferase reporter assays that measure replication of the viral genome (Table 2A). On the VEEV alphavirus (*R*)-**1** demonstrated potent activity while the (*S*)-**1** enantiomer was inactive. The enantioselectivity mirrored the profile of the CHIKV alphavirus (Table 1) and confirmed the broad-spectrum antiviral activity on both arthritic and encephalitic members of the family. No inhibition of Dengue virus (DENV) and West Nile virus (WNV), members of the Flavivirus genus of single-stranded positive-sense RNA viruses, and Ebola a single-stranded, negative-sense RNA virus that belongs to the Filoviridae family was observed at concentrations between 10–30 μM. The results demonstrated at least 100-fold selectivity of (*R*)-**1** for inhibition of alphaviruses over these other viruses.

**Table 2.** Selectivity over other RNA viruses.

**A. Luciferase reporter assays<sup>a</sup>**
| Virus | Family | Host Cell | (*R*)-1<br>EC<sub>50</sub> (μM) | (*S*)-1<br>EC<sub>50</sub> (μM) |
| --- | --- | --- | --- | --- |
| VEEV | Alpha | MRC5 | 0.27 | >40 |
| DENV2 | Flavi | Vero | >10 | >10 |
| DENV4 | Flavi | Vero | >10 | >10 |
| WNV | Flavi | Vero | >10 | >10 |
| Ebola | Filo | Vero | >30 | >30 |

<sup>a</sup>Assays performed over a 10-point dose range from 0.01–40 μM (VEEV), 0.001–10 μM (DENV2, DENV4, and WNV), and 0.01–30 μM (Ebola).

**B. CPE assays<sup>a</sup>**
| Virus | Family | Host Cell | (*R*)-1<br>EC<sub>50</sub> (μM) | (*S*)-1<br>EC<sub>50</sub> (μM) |
| --- | --- | --- | --- | --- |
| CHIKV | Alpha | Vero | 3.6 | >32 |
| WEEV | Alpha | Vero | 6.3 | >32 |
| YFV | Flavi | HuH7 | >32 | >32 |
| POWV | Flavi | BHK | >32 | >32 |
| JEV | Flavi | Vero | >32 | >32 |
| RSV | Pneumo | RD | >32 | >32 |
| ECHO-11 | Picorna | RD | >32 | >32 |
| ENTV68 | Picorna | RD | >32 | >32 |
| FluA H5N1 | Orthomyxo | MDCK | >32 | >32 |

<sup>a</sup>Assays performed over an 8-point dose range from 0.01–32 μM.

As an additional demonstration of antiviral selectivity, (*R*)-**1** and (*S*)-**1** were submitted for testing on a panel of RNA virus available through NIH Preclinical Services.^15^ These assays measured the cytopathic effect (CPE) effect of the viruses resulting from cell death (Table 2B). Across the nine RNA viruses, enantioselective activity of (*R*)-**1** but not (*S*)-**1** was seen against the alphaviruses CHIKV and WEEV. The lower potency of (*R*)-**1** on CHIKV in the CPE assay compared to the viral genome replication assay suggests that the latter is a more direct measure of the effect of nsP2hel inhibition. Importantly, no inhibition of the flavi-, pneumo-, picorna-, or orthomyxoviruses was seen with either enantiomer at concentrations >30 μM, further demonstrating the selectivity of (*R*)-**1** for the alphavirus family.

To assess selectivity against another viral RNA helicase, (*R*)-**1** was tested for inhibition of ATPase activity on SARS-CoV-2 NSP13, an SF1 helicase and the closest relative by primary sequence to alphavirus nsP2hel.^16^ (*R*)-**1** and (*S*)-**1** were also tested for inhibition of ATPase activity on four human DEAD-box family RNA helicases: MDA5, DDX1, DDX3X, and LGP2.^17^ In each case (Table 3), either no evidence of inhibition or only weak inhibition at high micromolar concentrations was observed with no enantioselectivity between (*R*)-**1** and (*S*)-**1**. Thus, (*R*)-**1** was >100-fold selective for CHIKV nsP2hel over five other RNA helicases.

**Table 3.**
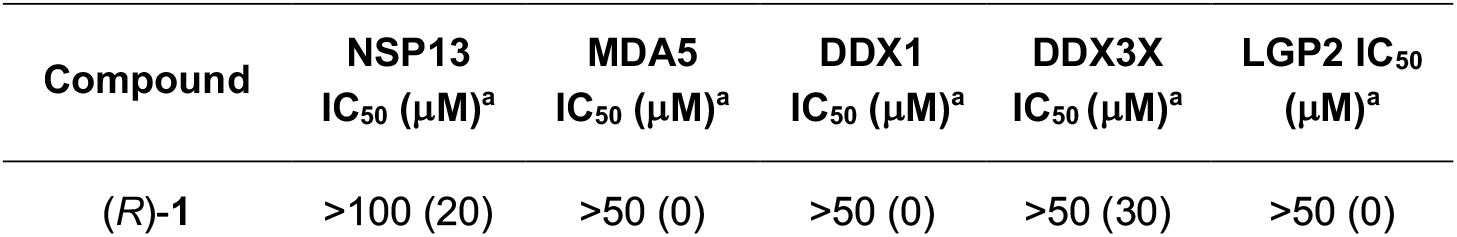
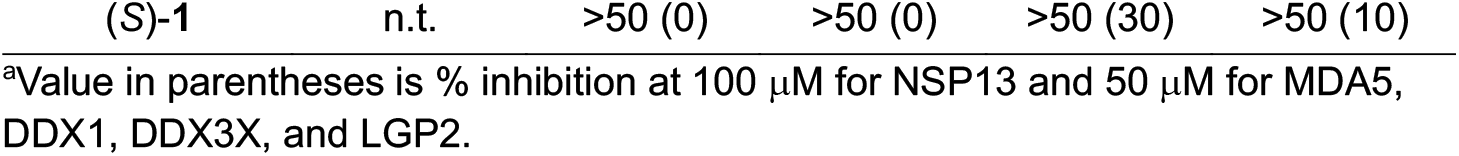
Selectivity over other helicases.

| Compound | NSP13<br>IC<sub>50</sub> (μM)<sup>a</sup> | MDA5<br>IC<sub>50</sub> (μM)<sup>a</sup> | DDX1<br>IC<sub>50</sub> (μM)<sup>a</sup> | DDX3X<br>IC<sub>50</sub> (μM)<sup>a</sup> | LGP2 IC<sub>50</sub><br>(μM)<sup>a</sup> |
| --- | --- | --- | --- | --- | --- |
| (*R*)-1 | >100 (20) | >50 (0) | >50 (0) | >50 (30) | >50 (0) |

| (S)-**1** | n.t. | >50 (0) | >50 (0) | >50 (30) | >50 (10) |
| --- | --- | --- | --- | --- | --- |

<sup>a</sup>Value in parentheses is % inhibition at 100 μM for NSP13 and 50 μM for MDA5, DDX1, DDX3X, and LGP2.

## Conclusion

This work establishes the (*R*)-enantiomer of oxaspiropiperidine **1**, designated RA-NSP2-1, as a potent and selective chemical probe for CHIKV nsP2hel. The resolution of racemic **1** revealed a striking >100-fold difference in antiviral activity and enzyme inhibition between (*R*)-**1** and its stereoisomer (*S*)-**1** (RA-NSP2-1N), underscoring the enantioselective mechanism of action. These findings were supported by matched potency differences in viral replication assays, ATPase inhibition, RNA unwinding assays, and ^19^F NMR target engagement studies, confirming that (*R*)-**1** directly binds to nsP2hel and inhibits its motor enzyme functions. The biological activity of (*R*)-**1** is highly specific to alphavirus nsP2, with no significant activity against other RNA viruses from other families, and minimal inhibition of human DEAD-box helicases or SARS-CoV-2 NSP13 helicase. This target selectivity, combined with potent cellular and biochemical activity, confirmed (*R*)-**1** (RA-NSP2-1) as a high-quality chemical probe suitable for dissecting nsP2hel function in cellular and biochemical settings.

Interestingly, NMR spectroscopy suggests that (*R*)-**1** exhibits conformational polymorphism due to amide rotamers and thermal mobility in the piperidine ring. This structural flexibility likely contributes to the resistance of (*R*)-**1** to crystallization but may also be functionally relevant to its action as a helicase inhibitor. Notably, only analogs of **1** with broad ^13^C NMR signals (indicating motion in the spirocyclic core) previously showed potent antiviral activity,^12^ suggesting a potential mechanistic link between thermal mobility of the ligand and allosteric inhibition. Such a property may enable (*R*)-**1** to accommodate dynamic conformational states of the RNA helicase, a feature that is often difficult to achieve with rigid, orthosteric inhibitors.^18^

The selective inhibition of nsP2hel by (*R*)-**1 (**RA-NSP2-1) and the availability of its inactive (*S*)-**1** (RA-NSP2-1N) counterpart position the compound pair as powerful tools to probe the dynamic motor function of a viral RNA helicase. In addition, these chemical probes may allow exploration of conformational adaptability as a determinant of efficacy for helicase inhibition. Importantly, (*R*)-**1 (**RA-NSP2-1) and (*S*)-**1** (RA-NSP2-1N) highlight the potential of enantioselective chemical probes to uncover molecular regulation of the alphavirus replication complex.

## Methods

### General Information

All reactions were conducted in oven-dried glassware under a dry nitrogen atmosphere unless otherwise specified. All reagents and solvents were obtained from commercial sources and used without further purification. No unexpected safety hazards were encountered during the syntheses. Analytical thin layer chromatography (TLC) was performed on pre-coated silica gel plates (200 μm, F_254_ indicator), visualized under UV light or by staining with iodine and KMnO_4_. Column chromatography utilized pre-loaded silica gel cartridges on a Biotage automated purification system. ^1^H and ^13^C NMR spectra were recorded in DMSO-*d*_6_, CD_3_CN and CD_3_OD at 400/500 and 101/126 MHz, respectively, on Bruker spectrometers. Chemical shifts (*δ*) are reported in parts per million (ppm) downfield from tetramethylsilane for ^1^H NMR, with major peaks designated as s (singlet), d (doublet), t (triplet), q (quartet), and m (multiplet), dd (doublet of doublets), td (triplet of doublets), qd (quartet of doublets), tt (triplet of triplets) and ddd (doublet of doublet of doublets). HRMS samples were analyzed at the UNC Department of Chemistry Mass Spectrometry Core Laboratory with a Q Exactive HF-X mass spectrometer. Analytical HPLC data were recorded on a Waters e2695 HPLC with PDA detector using a YMC Triart C18 (150 × 4.6 mm, 5 μm) column or Agilent 1260 Infinity II series with PDA detector using an EC-C18, 100 mm x 4.6 mm, 3.5 μm) column, with eluent 10−90% CH_3_CN in water at a flow rate of 1 mL/min. Enantiomeric ratio of the chiral compound was analyzed either on an Agilent 1260 infinity series HPLC with PDA detector using a Chiralpak IC (250 × 4.6 mm, 5 μm) column at a flow rate of 1.0 mL/min; mobile phase (A) 0.1% NH_3_ in hexane and (B) 0.1% NH_3_ in IPA:MeCN (70:30) with gradient condition 80:20 at 0.01 min, 50:50 at 5 min, 30:70 at 10 min up to 20 min, 80:20 at 20 to 25 min, or on a Waters SFC investigator with PDA detector using a Chiralpak AD-H (250 × 4.6 mm, 5 μm) column at a flow rate of 4.0 ml /min at 100 bar; mobile phase (A) liquid CO_2_ and (B) 0.1% NH_3_ in IPA:MeCN (50:50) with gradient condition 5% to 50% B over 5 min, then 50% B for up to 20 min. Optical rotations were measured with an Anton Paar MCP 200 Research or Autopol IV Automatic Polarimeter from Rudolph using a 10 cm microcell. (4-((1,1-Dioxido-1,2-thiazinan-2-yl)methyl)-1-oxa-9-azaspiro[5.5]undecan-9-yl)(1-phenylcyclopentyl)methanone (**1**) and tert-butyl 4-(aminomethyl)-1-oxa-9-azaspiro[5.5]undecane-9-carboxylate (**2**) were synthesized as previously described.^12^ All compounds were >95% Pure by HPLC analysis

### Chemistry

#### Chiral SFC separation of racemic (±)-1

Chiral SFC separation performed using a Shimadzu LC-20AP preparative pump with UV-detector set at 220 nm with mobile phase: (A) liquid CO_2_, (B) 0.1% NH_3_ in IPA:MeCN (50:50) using a Chiralpak AD-H column (250 × 30 mm, particle size 5 μm) and flow rate of 100 mL/min. *(±)-****1*** (100 mg, 0.211 mmol) yielded (*R*)-(4-((1,1-dioxido-1,2-thiazinan-2-yl)methyl)-1-oxa-9-azaspiro[5.5]undecan-9-yl)(1-phenylcyclopentyl)methanone (*R*)-**1** (40 mg, 40% yield): Rt 5.00 min; [α]_D_^25^ + 4.8 deg cm^3^ g^−1^ dm^−1^(c 0.0025 g cm^−3^, MeOH) and (*S*)-(4-((1,1-dioxido-1,2-thiazinan-2-yl)methyl)-1-oxa-9-azaspiro[5.5]undecan-9-yl)(1-phenylcyclopentyl)methanone (*S*)-**1** (26 mg, 26% yield): Rt 5.84 min; [α]_D_^25^ -4.4 deg cm^3^ g^−1^ dm^−1^(c 0.0025 g cm^−3^, MeOH).

#### tert-Butyl 4-(aminomethyl)-1-oxa-9-azaspiro[5.5]undecane-9-carboxylate D-DTTA salt (R- 2)_2_•D-DTTA

To a stirred solution of tert-butyl 4-(aminomethyl)-1-oxa-9-azaspiro[5.5]undecane-9-carboxylate (**2**) (10 aliquots of 500 mg, 1.760 mmol, 1.0 eq) in EtOH (5 volumes) at RT was added di-p-toluoyl-D-tartaric acid (339.7 mg, 0.8802 mmol, 0.5 eq). to each aliquot. The resulting mixtures were stirred at 90 °C for 30 min until the solution became clear. The resulting solutions were allowed to cool to RT resulting in the precipitation of a solid. The solid from each of the aliquots was collected by filtration, washed twice with EtOH, and dried under reduced pressure to afford a first crop of 6.7 g of D-DTTA salt with ∼90% ee (40% total yield, 80% yield of *R*-1 enantiomer). The D-DTTA salt was recrystallized twice using ethanol (5 volumes) to obtain (*R*)-tert-butyl 4-(aminomethyl)-1-oxa-9-azaspiro[5.5]undecane-9-carboxylate D-DTTA salt (*R*-**2**)_2_*•D-* DTTA as a white solid (1.6 g, 10% yield). m.p.: 182°C–184°C; ^1^H NMR (500 MHz, CD_3_OD) δ 8.05 (d, *J* = 8.2 Hz, 4H), 7.28 (d, *J* = 7.8 Hz, 4H), 5.81 (s, 2H), 3.77–3.65 (m, 6H), 3.60 (td, *J* = 12.2, 2.4 Hz, 2H), 3.18–3.16 (m, 2H), 2.99 (br s, 2H), 2.70 (dd, *J* = 7.0, 1.7 Hz, 4H), 2.41 (s, 6H), 2.19– 2.11 (m, 2H), 2.08–1.99 (m, 2H), 1.65–1.62 (m, 2H), 1.59–1.55 (m, 2H), 1.53–1.46 (m, 4H), 1.45 (s, 18H), 1.20–1.11 (m, 2H), 1.15 (qd, *J* = 12.6, 5.2 Hz, 2H), 1.00 (t, *J* = 12.8 Hz, 2H); ^13^C NMR (126 MHz, CD_3_OD) δ 174.0, 167.9, 156.6, 144.9, 131.2, 130.0, 129.3, 81.0, 76.9, 71.3, 60.9, 46.1, 40.5, 40.1, 30.9, 30.2, 30.0, 28.7, 21.7; [α]_D_^25^ +77.81 deg cm^3^ mol^−1^ dm^−1^(c 0.1 M, MeOH).

#### tert-Butyl 4-(aminomethyl)-1-oxa-9-azaspiro[5.5]undecane-9-carboxylate L-DTTA salt (S- 2)2 DTTA). (S-2)_2_• L-DTTA was synthesized by reacting L-DTTA with the racemic 2: [α]_D_^25^ -60 deg cm^3^ mol^−1^ dm^−1^ (c 0.1M, MeOH)

#### tert-Butyl (R)-4-((4-methoxybenzamido)methyl)-1-oxa-9-azaspiro[5.5]undecane-9-carboxylate (R)-3

A solution of tert-butyl 4-(aminomethyl)-1-oxa-9-azaspiro[5.5]undecane-9-carboxylate D-DTTA salt *(R-****2***)_*2*_*•D-DTTA* (250 mg, 1.0 eq) in 1M KOH solution (10 volumes) at RT was stirred for 30 min at RT. The resulting mixture was extracted with 10% MeOH in DCM (25 ml x 3). The organic layers were combined and dried over sodium sulphate, filtered and concentrated to afford tert-butyl (*R*)-4-(aminomethyl)-1-oxa-9-azaspiro[5.5]undecane-9-carboxylate (*R*)-**2** as a colorless gum (0.070 g, 94% yield). LCMS: m/z [M-56]^+^ 229. The crude product was used directly in the next step without further purification. To a stirred solution of tert-butyl (*R*)-4-(aminomethyl)-1-oxa-9-azaspiro[5.5]undecane-9-carboxylate (0.070 g, 1.0 eq) in DCM (10 volumes) was added Et_3_N (0.1 mL, 3.0 eq) and the mixture stirred for 10 min at RT. The resulting mixture was cooled to 0 °C and 4-methoxybenzoyl chloride (0.050 g, 1.2 eq) was added dropwise and then stirred at RT for 1 hour. The mixture was quenched with ice-cold water (60 mL) and extracted with EtOAc (2 × 25 mL). The combined organic layers were washed with brine, dried over Na_2_SO_4_, filtered, and concentrated to obtain the crude product which was purified by CombiFlash, eluting with 67% ethyl acetate in hexane. tert-Butyl (*R*)-4-((4-methoxybenzamido)methyl)-1-oxa-9-azaspiro[5.5]undecane-9-carboxylate (*R*)-**3** was isolated as a colorless sticky gum (0.069 g, 67% yield): ^1^H NMR (400 MHz, DMSO-d_6_) δ 8.30 (s, 1H), 7.81 (d, *J* = 6.8 Hz, 2H), 6.97 (d, *J* = 8.8 Hz, 2H), 3.80–3.79 (m, 3H), 3.66–3.60 (m, 3H), 3.57–3.48 (m, 1H), 3.08 (br s, 2H), 2.99 (br s, 1H), 2.09–2.05 (m, 1H), 1.99–1.95 (m, 2H), 1.58–1.55 (m, 2H), 1.37 (s, 9H), 1.19–1.15 (m, 3H), 1.10– 1.07 (m, 1H), 0.98–0.91 (m, 1H); LCMS m/z [M-100] ^+^ 319.2; chiral HPLC 98.3 %; [α]_D_^25^ + 9.4 deg cm^3^ mol^−1^ dm^−1^(c 0.1M, MeOH).

#### tert-Butyl (S)-4-((4-methoxybenzamido)methyl)-1-oxa-9-azaspiro[5.5]undecane-9-carboxylate (S)-3

(*S)*-**3** was synthesized using the same procedure from tert-butyl 4-(aminomethyl)-1-oxa-9-azaspiro[5.5]undecane-9-carboxylate L-DTTA salt (*S*-**2**)_2_.•L-DTTA: chiral HPLC 98.4% ee; [α]_D_^25^ -7 deg cm^3^ mol^−1^ dm^−1^(c 0.1M, MeOH).

(*R*)-(4-((1,1-dioxido-1,2-thiazinan-2-yl)methyl)-1-oxa-9-azaspiro[5.5]undecan-9-yl)(1-phenylcyclopentyl)methanone (*R*-(+)-**1**). To a stirred solution of tert-butyl (*R*)-4-(aminomethyl)-1-oxa-9-azaspiro[5.5]undecane-9-carboxylate (*R*)-**2** (0.610 g, 1.0 eq) in DMF (10 mL) at RT was added 4-chloro-1-butylsulfonyl chloride (0.492 g, 1.2 eq) and triethylamine (TEA) (0.9 mL, 3.0 eq). The reaction mixture was stirred at RT for 3 h. Upon completion of the reaction, the mixture was quenched with ice-cold water (150 mL) and extracted with EtOAc (75 mL × 3). The combined organic layers were washed with brine, dried over Na_2_SO_4_, filtered, and concentrated to obtain the crude product which was purified by CombiFlash, eluting with 60% EtOAc in hexane. tert-Butyl (*R*)-4-(((4-chlorobutyl)sulfonamido)methyl)-1-oxa-9-azaspiro[5.5]undecane-9-carboxylate was isolated as a colorless gum (0.910 g, 97% yield): LCMS-ESI+ (m/z) [M-56] ^+^= 383.3. To a stirred solution of tert-butyl (*R*)-4-(((4-chlorobutyl)sulfonamido)methyl)-1-oxa-9-azaspiro[5.5]undecane-9-carboxylate (0.900 g, 1.0 eq.) in MeCN (10 volumes) at RT was added DBU (1.52 mL, 5.0 eq.). The mixture was stirred at 70 °C for 2 h, then diluted with water (150 mL) and extracted with EtOAc (75 ml x 3). The combined organic layers were washed with brine, dried over Na_2_SO_4_, filtered, and concentrated to obtain the crude product which was purified by CombiFlash, eluting with 50% EtOAc in hexane to afford tert-butyl (*R*)-4-((1,1-dioxido-1,2-thiazinan-2-yl)methyl)-1-oxa-9-azaspiro[5.5]undecane-9-carboxylate as a white foam (0.630 g, 74% yield): LCMS-ESI+ (m/z) [M-100]^+^= 303.1. To a stirred solution of tert-butyl (*R*)-4-((1,1-dioxido-1,2-thiazinan-2-yl)methyl)-1-oxa-9-azaspiro[5.5]undecane-9-carboxylate (0.620 g, 1.0 eq.) in DCM (10 volumes) at RT was added 4M HCl in dixoane (2 mL). The mixture was stirred at RT for 2 h then evaporated and triturated with Et_2_O to afford (*R*)-2-((1-oxa-9-azaspiro[5.5]undecan-4-yl)methyl)-1,2-thiazinane 1,1-dioxide hydrochloride as a white solid (0.515 g, 99% yield): LCMS-ESI+ (m/z) 303.1. To a stirred solution of (*R*)-2-((1-oxa-9-azaspiro[5.5]undecan-4-yl)methyl)-1,2-thiazinane 1,1-dioxide hydrochloride (0.50 g, 1.2 eq.) and 1-phenyl-1-cyclopentanecarboxylic acid (0.235 g, 1.0 eq.) in DMF (10 mL) at RT was added EDC•HCl (0.355 g, 1.5 eq), HOBt (0.151 g, 0.8 eq), and DIPEA (1.08 mL, 5.0 eq). The mixture was stirred at RT for 6 h, then diluted with water (150 mL) and extracted with EtOAc (75 mL × 3). The combined organic layers were washed with brine, dried over Na_2_SO_4_, filtered, and concentrated to obtain the crude product which was purified by CombiFlash, eluting with 50% EtOAc in hexane. Acetonitrile (2 mL) and water (2 mL) were then added to the white foam and following lyophilization (*R*)-(4-((1,1-dioxido-1,2-thiazinan-2-yl)methyl)-1-oxa-9-azaspiro[5.5]undecan-9-yl)(1-phenylcyclopentyl)methanone (*R*)-(+)-**1** was isolated as a white amorphous powder (0.33 g, 56% yield): ^1^H NMR (500 MHz, CD_3_CN): δ 7.35–7.29 (m, 2H), 7.25– 7.18 (m, 3H), 4.15 (br s, 1H), 3.60 (dd, *J* = 12.1, 4.9 Hz, 1H), 3.54–3.34 (m, 1H), 3.31–3.18 (m, 3H), 3.08–2.95 (m, 1H), 2.94–2.90 (m, 2H), 2.88–2.73 (m, 3H), 2.50–2.26 (m, 2H), 2.11–2.02 (m, 3H), 1.88–1.62 (m, 6H), 1.57–1.52 (m, 3H), 1.44–1.40 (m, 2H), 1.31–1.16 (m, 1H), 1.02 (qd, *J* = 12.6, 5.1 Hz, 1H), 0.93–0.62 (m, 3H); ^13^C NMR (126 MHz, CD_3_CN): δ 174.5, 147.2, 129.6, 126.9, 126.0, 71.0, 60.8, 59.3, 54.1, 51.2, 48.0, 40.7, 31.2, 30.7, 26.1, 24.6, 20.8; HRMS (ESI) *m/z* [M + H]^+^ calculated for C_26_H_39_N_2_O_4_S 475.2630, found 475.2622; chiral HPLC > 99% ee; [α]_D_^25^ + 4.40 deg cm^3^ g^−1^ dm^−1^(c 0.0025 g cm^−3^, MeOH).

#### (S)-(4-((1,1-dioxido-1,2-thiazinan-2-yl)methyl)-1-oxa-9-azaspiro[5.5]undecan-9-yl)(1-phenylcyclopentyl)methanone (S-(-)-1)

*S*-(-)-**1** was synthesized using the same procedure tert-butyl 4-(aminomethyl)-1-oxa-9-azaspiro[5.5]undecane-9-carboxylate L-DTTA salt (*S*-**2**)_2_.•L-DTTA: HRMS (ESI) *m/z* [M + H]^+^ calculated for C_26_H_39_N_2_O_4_S 475.2630, found 475.2623; chiral HPLC > 99% ee; [α]_D_^25^–4.4 deg cm^3^ g^−1^ dm^−1^(c 0.0025 g cm^−3^, MeOH).

#### (R)-(1-(3,5-difluorophenyl)cyclopentyl)(4-((1,1-dioxido-1,2-thiazinan-2-yl)methyl)-1-oxa-9-azaspiro[5.5]undecan-9-yl)methanone (R)-4

To a stirred solution of 1-(3,5-difluorophenyl)cyclopentane-1-carboxylic acid (58 mg, 1.1 eq, 0.25 mmol) in DMF (1 mL) at 0 °C was added HATU (0.13 g, 0.35 mmol, 1.5 eq) and DIPEA (0.19 mL, 0.93 mmol, 5 eq). After 10 min, (*R*)-2-((1-oxa-9-azaspiro[5.5]undecan-4-yl)methyl)-1,2-thiazinane 1,1-dioxide hydrochloride (75 mg, 0.22 mmol, 1 eq) was added. The mixture was stirred at RT for 16 h, then diluted with water and extracted with EtOAc. The organic extract was washed with brine, dried over Na_2_SO_4_, filtered and concentrated to give the crude product which was purified by CombiFlash, eluting with 60% EtOAc in hexane to afforded (*R*)-(1-(3,5-difluorophenyl)cyclopentyl)(4-((1,1-dioxido-1,2-thiazinan-2-yl)methyl)-1-oxa-9-azaspiro[5.5]undecan-9-yl)methanone (*R*)-**4** (68 mg, 0.14 mmol, 59 %) as a white foam: ^1^H NMR (500 MHz, DMSO): δ 7.12–7.08 (m, 1H), 6.87–6.83 (m, 2H), 4.13–4.00 (m, 1H), 3.61–3.58 (m, 1H), 3.45–3.42 (m, 1H), 3.27–3.25 (t, *J* = 5.5 Hz, 2H), 3.17– 3.08 (m, 1H), 3.05–2.89 (m, 3H), 2.83–2.73 (m, 3H), 2.37–2.31 (m, 2H), 2.18–1.95 (m, 3H), 1.88– 1.78 (m, 3H), 1.69–1.48 (m, 7H), 1.44–1.30 (m, 2H), 1.27–1.05 (m, 1H), 1.03–0.74 (m, 3H); ^13^C NMR (126 MHz, DMSO) δ 171.9, 162.6 (dd, *J* = 247, 12.6 Hz), 150.4, 108.2 (d, *J* = 25 Hz), 101.7 (t, *J* = 25), 69.7, 59.6, 58.0, 52.8, 50.0, 46.6, 30.1, 29.325.0, 24.9, 23.0, 19.5; HRMS (ESI) *m/z* [M + H]^+^ calculated for C_26_H_37_F_2_N_2_O_4_S 511.2442, found 511.2430; [α]_D_^25^ +2.50 deg cm^3^ g^−1^ dm^−1^(c 0.0020 g cm^−3^, MeOH).

#### (S)-(1-(3,5-difluorophenyl)cyclopentyl)(4-((1,1-dioxido-1,2-thiazinan-2-yl)methyl)-1-oxa-9-azaspiro[5.5]undecan-9-yl)methanone (S)-4

(*S*)-**4** was synthesized using the same procedure from (*S*)-2-((1-oxa-9-azaspiro[5.5]undecan-4-yl)methyl)-1,2-thiazinane 1,1-dioxide: [α]_D_^25^ -2.00 deg cm^3^ g^−1^ dm^−1^(c 0.0020 g cm^−3^, MeOH).

### X-ray Crystallography

A colorless crystal of (*R*)-**2** grown from MeOH (approximate dimensions 0.210 × 0.070 × 0.020 mm^3^) was placed onto the tip of MiTeGen and mounted on a Bruker D8 VENTURE diffractometer and measured at 150 K.

#### Data collection

A preliminary set of cell constants was calculated from reflections harvested from a set of 180 frames to produce orientation matrices determined from 483 reflections. The data collection was carried out using Cu Kα radiation (graphite monochromator) with theta-dependent frame window of 1–7 sec and a detector distance of 3.7 cm. A randomly oriented region of reciprocal space was surveyed to achieve complete data with a redundancy of 5.9. Sections of frames were collected with 1.0º steps in ω and ϕ scans. Data to a resolution of 0.82 Å were considered in the reduction. Final cell constants were calculated from the xyz centroids of 9838 strong reflections from the actual data collection after integration (SAINT).^19^ The intensity data were corrected for absorption (SADABS).^20^ Additional crystal and refinement information is listed in Table S1.

#### Structure solution and refinement

Space group *C* 2 was determined based on intensity statistics and systematic absences. The structure was solved using SHELXT^21^ and refined (full-matrix-least squares) using the Oxford University Crystals for Windows system.^22^ The intrinsic-phasing solution provided most non-hydrogen atoms from the E-map. Full-matrix least squares and difference Fourier cycles were performed to locate the remaining non-hydrogen atoms. The structure (*R*-**2**)_2_•D-DTTA exhibits significant disorder on the Boc amide. Addition disorder originated from the piperidine ring and consisted of two components each exhibiting its own two-part positional disorder in the Boc amide. The disorder was modeled such that the occupancies of the major and minor components of piperidine summed to one, with the sub-disorder parts within each component summed to the respective component’s occupancy. All non-hydrogen atoms were refined using anisotropic displacement parameters. Hydrogen atoms bonded to nitrogen were obtained from the difference map and refined isotropically, while all other hydrogen atoms were placed in ideal positions and refined as riding atoms. The final full matrix least squares refinement converged to R1 = 0.0865 and wR2 = 0.2602 (F^2^, all data).

#### Structure description

The crystal structure was confirmed as the (R)-**2** amine and D-DTTA in a 2:1 ratio. The Flack parameter for chirality was determined to be 0.0(3). The less-than-ideal standard deviations (esds) were attributed to the predominance of light atoms in the structure. Crystallographic data has been deposited in the Cambridge Structural Database under CCDC 2034666. These data can be obtained free of charge from www.ccdc.cam.ac.uk/structures.

### *Ab Initio* Calculation

DFT calculations were performed using Gaussian package, version 16C01, with tight SCF convergence and ultrafine grids. The function employed was B3LYP with basis set Pople’s triple-zeta 6-311+G(d,p).^23^ The dispersion correction was included the calculations using the D3 version of Grimme’s dispersion with Becke-Johnson damping.^24^ A full structure optimization of (*R*-**2**)_2_•D-DTTA was performed starting with the X-ray crystal structure coordinate as the initial geometry. Basis set superposition error (BSSE)^25^ corrected calculations were used to calculate the interaction energy between the amine **2** and D-DTTA.

### Biological Evaluation

#### CHIKV antiviral and helicase assays

CHIKV viral replication, nsP2 ATPase, and nsP2 unwindase activity of test compounds were determined using the previously described protocols.^12^

#### Human RNA helicase assays

ATPase activity of MDA5, DDX1, LGP2, and DDX3X were monitored in vitro using a bioluminescent assay protocol as previously described.^26^

#### Viral genome replication assays

The effect of test compounds on viral replication was measured using luciferase assays as previously described for alphavirus VEEV,^12^ flaviviruses DENV and WNV,^27^ and Ebola virus.^28^ and enabled within the READDI-AViDD Center.^29^

^*19*^*F NMR*. Spectra were captured in the presence of CHIKV nsP2hel domain protein using the previously described protocol.^12^ Target engagement was determined by measurement of the peak area of the ^19^F resonance at -109.82 ppm in the absence and presence of nsP2hel protein.

## Supporting Information

Figure S1: ^1^H NMR spectrum (*R*-**2**)_2_•D-DTTA. Figure S2: Molecular structure of (*R*-**2**)_2_•D-DTTA. Figure S3: Four conformers of (*R*-**2**)_2_•D-DTTA seen in the SC-XRD structure. Figure S4: Energy-minimized structures of (*R*-**2**)_2_•D-DTTA. Figure S5: Enantioselectivity of **4**. Figures S6– S24 ^1^H NMR, ^13^C NMR, chiral HPLC, and analytical HPLC spectra. Table S1–S6: X-ray crystallographic refinement data.

## Acknowledgement

The Structural Genomics Consortium (SGC) is a registered charity (no: 1097737) that receives funds from Bayer AG, Boehringer Ingelheim, Bristol Myers Squibb, Genentech, Genome Canada, through Ontario Genomics Institute [OGI-196], EU/EFPIA/OICR/McGill/KTH/Diamond Innovative Medicines Initiative 2 Joint Under-taking [EUbOPEN grant 875510], Janssen, Merck KGaA (also known as EMD in Canada and the US), Pfizer, and Takeda. The research reported in this publication was supported by NIH grant 1U19AI171292-01 (READDI-AViDD Center) and in part by the NC Biotech Center Institutional Support Grant 2018-IDG-1030 and by NIH grant S10OD032476 for upgrading the 500 MHz NMR spectrometer in the UNC Eshelman School of Pharmacy NMR Facility. X-ray crystallography was supported by the National Science Foundation under Grant No. (CHE-2117287). This project was also supported by equipment purchased with support by the North Carolina State Policy Collaboratory at UNC Chapel Hill, as well as equipment purchased with support by the Rapidly Emerging Antiviral Drug Development Initiative at UNC Chapel Hill with funding from the North Carolina Coronavirus State and Local Fiscal Recovery Funds program, appropriated by the North Carolina General Assembly. We thank Ashish Pathak for coordinating RNA virus selectivity testing through NIH preclinical services and Piramal Pharma Solution for additional chemistry support.

## Abbreviations

BSSE: Basis set superposition error
CHIKV: Chikungunya virus
CPE: Cytopathic effect DENV Dengue virus
DTTA: Di-*p*-toluoyl tartaric acid
EEEV: Eastern equine encephalitis virus
nsP2hel: Non-structural protein 2 RNA helicase
MTL: Methyl transferase-like
SFC: Supercritical fluid chromatography
WNV: West Nile virus
SC-XRD: Single crystal X-ray diffraction

## Table of Contents Graphic

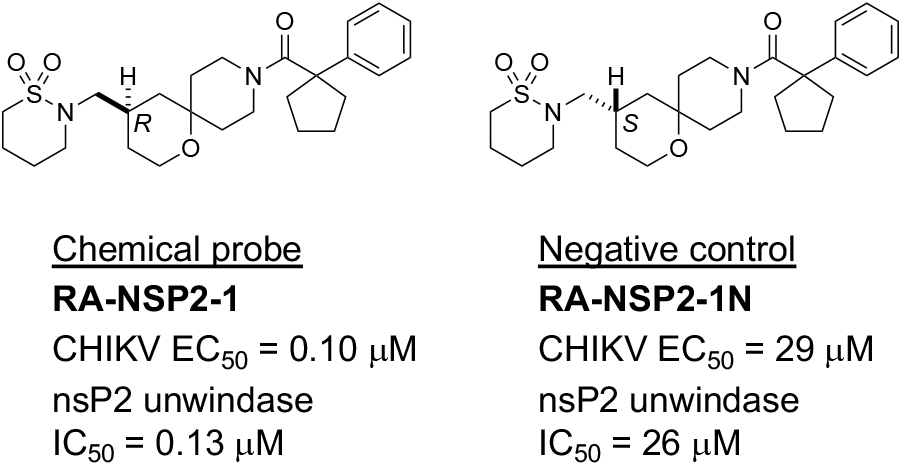

## References

(1) Skidmore, A. M.; Bradfute, S. B. The life cycle of the alphaviruses: From an antiviral perspective. Antiviral Res 2023, 209, 105476. doi: 10.1016/j.antiviral.2022.105476.

(2) Morens, D. M.; Folkers, G. K.; Fauci, A. S. Eastern Equine Encephalitis Virus - Another Emergent Arbovirus in the United States. N Engl J Med 2019, 381 (21), 1989–1992. doi: 10.1056/NEJMp1914328.

(3) Massachusetts arbovirus update. Commonwealth of Massachusetts, https://www.mass.gov/info-details/massachusetts-arbovirus-update (accessed September 1st, 2024).

(4) Croddy, E. The Post-World War II Era and the Korean War. In Chemical and Biological Warfare: A Comprehensive Survey for the Concerned Citizen, Springer, 2002; pp 30–31.

(5) Bohnsack, K. E.; Yi, S.; Venus, S.; Jankowsky, E.; Bohnsack, M. T. Cellular functions of eukaryotic RNA helicases and their links to human diseases. Nat Rev Mol Cell Biol 2023, 24 (10), 749–769. doi: 10.1038/s41580-023-00628-5.

(6) Lang, N.; Jagtap, P. K. A.; Hennig, J. Regulation and mechanisms of action of RNA helicases. RNA Biol 2024, 21 (1), 24–38. doi: 10.1080/15476286.2024.2415801.

(7) Fairman-Williams, M. E.; Guenther, U. P.; Jankowsky, E. SF1 and SF2 helicases: family matters. Curr Opin Struct Biol 2010, 20 (3), 313–324. doi: 10.1016/j.sbi.2010.03.011.

(8) Wang, S.; Mahalingam, S.; Merits, A. Alphavirus nsP2: A Multifunctional Regulator of Viral Replication and Promising Target for Anti-Alphavirus Therapies. Rev Med Virol 2025, 35 (2), e70030. doi: 10.1002/rmv.70030.

(9) Das, P. K.; Merits, A.; Lulla, A. Functional cross-talk between distant domains of chikungunya virus non-structural protein 2 is decisive for its RNA-modulating activity. J Biol Chem 2014, 289 (9), 5635–5653. doi: 10.1074/jbc.M113.503433.

(10) Law, Y. S.; Utt, A.; Tan, Y. B.; Zheng, J.; Wang, S.; Chen, M. W.; Griffin, P. R.; Merits, A.; Luo, D. Structural insights into RNA recognition by the Chikungunya virus nsP2 helicase. Proc Natl Acad Sci U S A 2019, 116 (19), 9558–9567. doi: 10.1073/pnas.1900656116.

(11) Tan, Y. B.; Chmielewski, D.; Law, M. C. Y.; Zhang, K.; He, Y.; Chen, M.; Jin, J.; Luo, D. Molecular architecture of the Chikungunya virus replication complex. Sci Adv 2022, 8 (48), eadd2536. doi: 10.1126/sciadv.add2536.

(12) Bose, M. R.; Sears, J. D.; Talbot, K. M.; Su, Y.-W. N.; Houliston, S.; Hossain, M. A.; Davis-Gilbert, Z. W.; Zhao, C.; Oh, H. J.; Brown, P. J.; et al. Identification of Direct-acting nsP2 Helicase Inhibitors with Anti-alphaviral Activity. bioRxiv 2025, 2025.2003.2004.641060. doi: 10.1101/2025.03.04.641060.

(13) Selvaratnam, L.; Willson, T. M.; Schapira, M. Structural Chemistry of Helicase Inhibition. J Med Chem 2025, 68 (4), 4022–4039. doi: 10.1021/acs.jmedchem.4c01909.

(14) Yu, L.; Reutzel-Edens, S. M.; Mitchell, C. A. Crystallization and Polymorphism of Conformationally Flexible Molecules: Problems, Patterns, and Strategies. Organic Process Research & Development 2000, 4 (5), 396–402. doi: 10.1021/op000028v.

(15) Preclinical and Clinical Resources for Antiviral Program for Pandemics Projects. National Insititute of Allergy and Infectious Diseases, https://www.niaid.nih.gov/research/antiviral-program-pandemics-resources (accessed April 17th, 2025).

(16) Selvaratnam, L. Multiple Sequence Alignment and Phylogenetic Tree of Human and Viral Helicases. Zenodo 2023. doi: 10.5281/zenodo.7569199.

(17) Andrisani, O.; Liu, Q.; Kehn, P.; Leitner, W. W.; Moon, K.; Vazquez-Maldonado, N.; Fingerman, I.; Gale, M., Jr. Biological functions of DEAD/DEAH-box RNA helicases in health and disease. Nat Immunol 2022, 23 (3), 354–357. doi: 10.1038/s41590-022-01149-7.

(18) Tee, W.-V.; Berezovsky, I. N. Allosteric drugs: New principles and design approaches. Current Opinion in Structural Biology 2024, 84, 102758. doi: 10.1016/j.sbi.2023.102758.

(19) SAINT, Bruker Analytical X-Ray Systems, Madison, WI.

(20) Blessing, R.H. An empirical correction for absorption anisotropy. Acta Cryst. 1995, A51, 33 – 38.

(21) Sheldrick, G. M. SHELXT – Integrated space-group and crystal-structure determination. Acta Cryst. 2015, 71, 3–8.

(22) P. W. Betteridge, J. R. C., R. I. Cooper, K. Prout, D. J. Watkin. CRYSTALS version 12: software for guided crystal structure analysis. J. Appl. Cryst. 2003, 36, 1487.

(23) Becke, A. D. Density-functional thermochemistry. III. The role of exact exchange. The Journal of Chemical Physics 1993, 98 (7), 5648–5652. doi: 10.1063/1.464913 (acccessed 4/22/2025).

(24) Grimme, S.; Ehrlich, S.; Goerigk, L. Effect of the damping function in dispersion corrected density functional theory. J Comput Chem 2011, 32 (7), 1456–1465. doi: 10.1002/jcc.21759.

(25) Boys, S. F.; and Bernardi, F. The calculation of small molecular interactions by the differences of separate total energies. Some procedures with reduced errors. Molecular Physics 1970, 19 (4), 553–566. doi: 10.1080/00268977000101561.

(26) Li, F.; Chan, U. H.; Perez, J. G.; Zeng, H.; Chau, I.; Li, Y.; Seitova, A.; Halabelian, L. ATPase activity profiling of three human DExD/H-box RNA helicases. SLAS Discov 2025, 32, 100229. doi: 10.1016/j.slasd.2025.100229.

(27) Baker, C.; Liu, Y.; Zou, J.; Muruato, A.; Xie, X.; Shi, P. Y. Identifying optimal capsid duplication length for the stability of reporter flaviviruses. Emerg Microbes Infect 2020, 9 (1), 2256–2265. doi: 10.1080/22221751.2020.1829994.

(28) Halfmann, P.; Kim, J. H.; Ebihara, H.; Noda, T.; Neumann, G.; Feldmann, H.; Kawaoka, Y. Generation of biologically contained Ebola viruses. Proc Natl Acad Sci U S A 2008, 105 (4), 1129–1133. doi: 10.1073/pnas.0708057105.

(29) Rapidly Emerging Antiviral Drug Development Initiative AViDD Center. The University of North Carolina, https://readdi-ac.org/ (accessed April 19th, 2025).

