## Supporting Information for "An Enantioselective Chemical Probe for Chikungunya nsP2 Helicase with Antialphaviral Activity"

Table of Contents

| | Description | Page |
|---|---|---|
| Figure S1 | $^1$H NMR spectrum (*R*-**2**)$_2$•D-DTTA | S3 |
| Figure S2 | Molecular structure of (*R*-**2**)$_2$•D-DTTA | S4 |
| Figure S3 | Four conformers of (*R*-**2**)$_2$•D-DTTA seen in the XRD structure | S5 |
| Figure S4 | Energy-minimized structures of (*R*-**2**)$_2$•D-DTTA | S6 |
| Figure S5 | Enantioselectivity of **4** | S6 |
| Figure S6–S24 | $^1$H NMR, 13C NMR, chiral HPLC, and analytical HPLC | S7-S16 |
| Table S1–S6 | X-ray crystallographic Refinement Data | S17–S29 |

**Figure S1.** $^1H$ NMR spectrum of (*R*-**2**)$_2$•D-DTTA

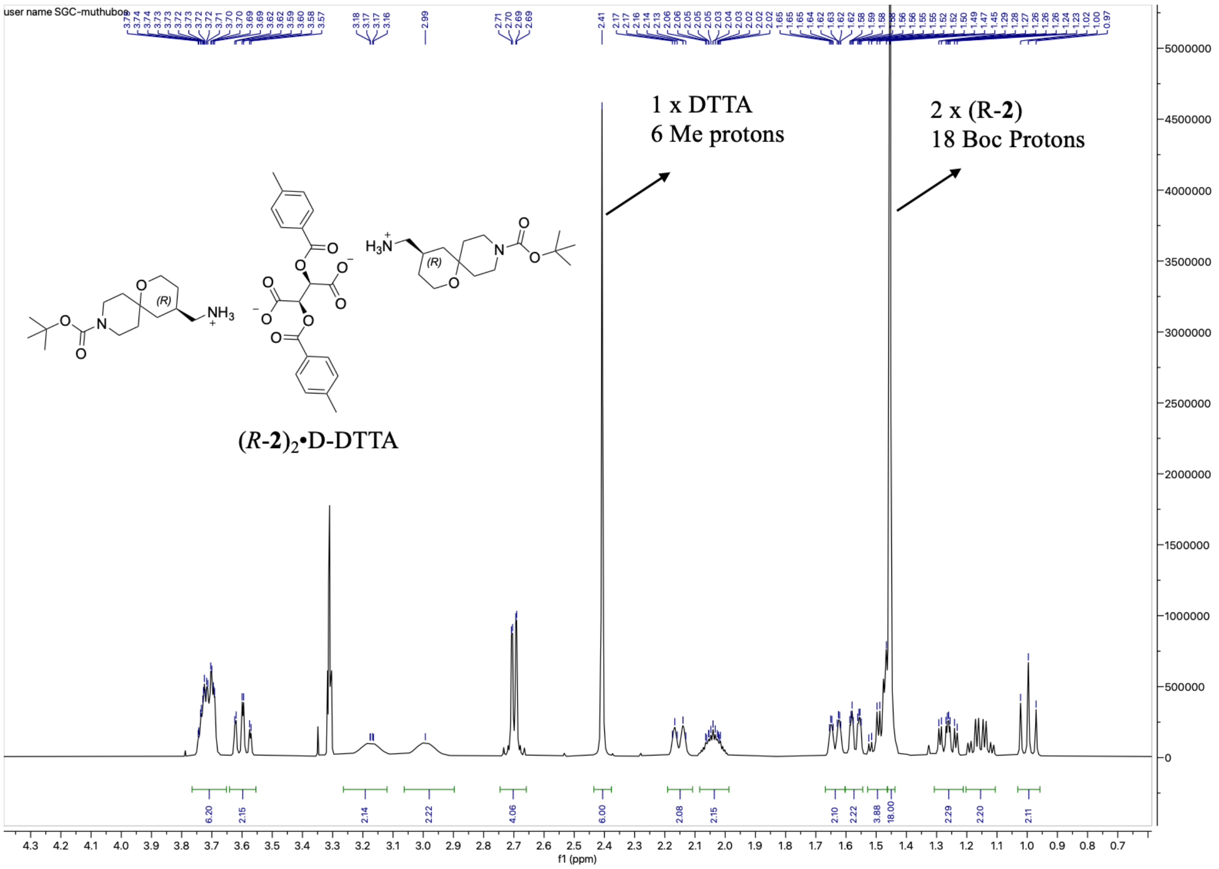

Integration of the Me protons from DTTA and $^t$Bu protons from (*R*)-**2** indicates a 2:1 ratio

**Figure S2.** Molecular structure of ($R$-**2**)$_2$•D-DTTA

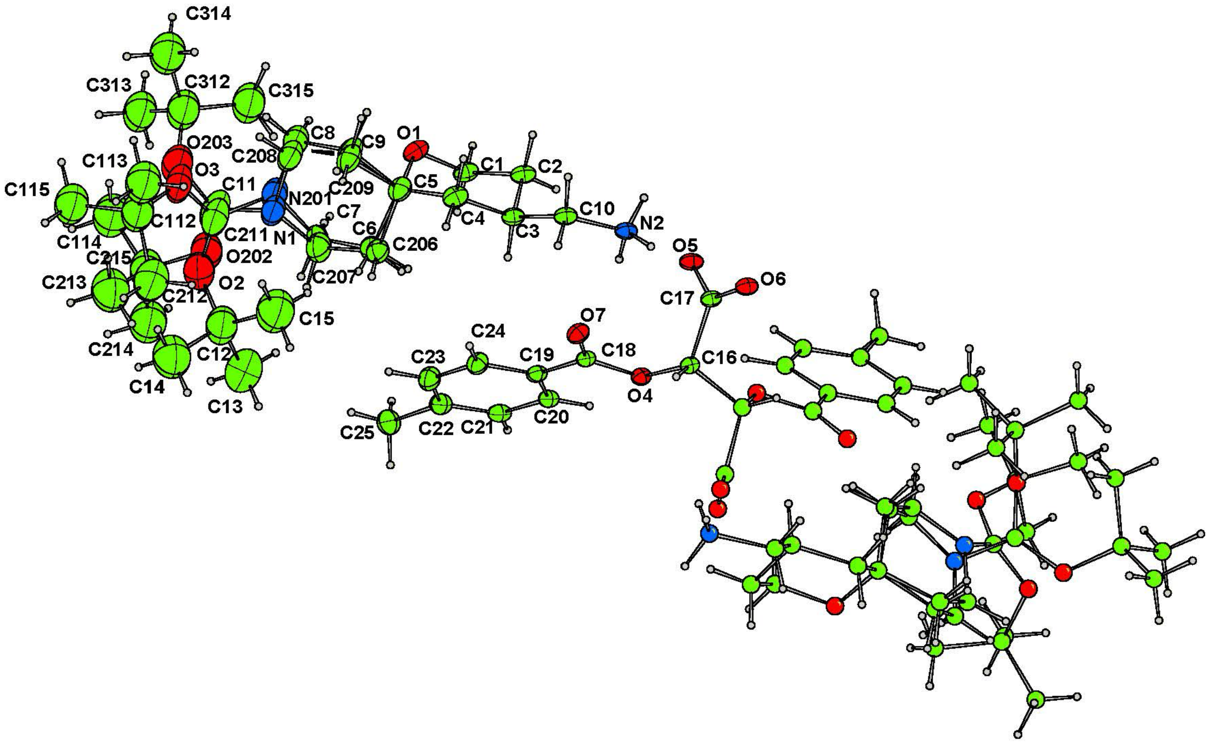

The asymmetric unit was labeled with atom numbers and displayed in ellipsoid view. The individual atom numbering in this view was used in the crystallographic refinement Tables S1–S6. Atom C3 in the tetrahydropyran ring of amine **2** has the *R* absolute configuration. This atom is C-4 of amine **2** using IUPAC numbering convention (c.f. Figure 2 of the main text).

**Figure S3.** Four conformers of ($R$-**2**)$_2$•D-DTTA seen in the XRD structure

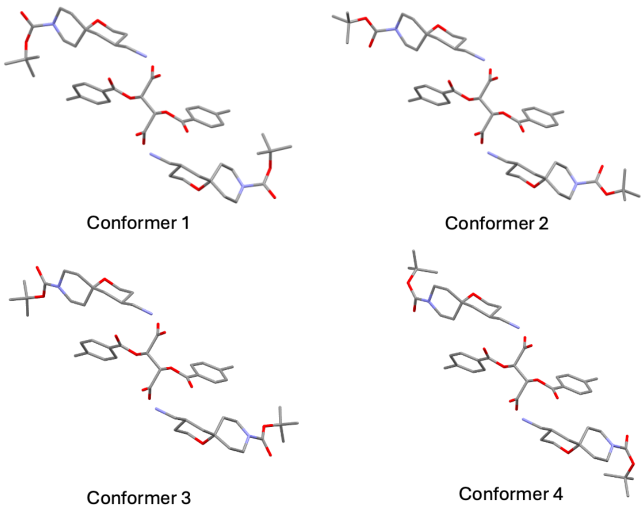

Conformers 1 and 2 are rotamers of the C-N bond with the first piperidine conformation. Conformers 3 and 4 are rotamers of the C-N bond with a second piperidine conformation.

**Figure S4.** Energy-minimized structures of ($R$-**2**)$_2$•D-DTTA

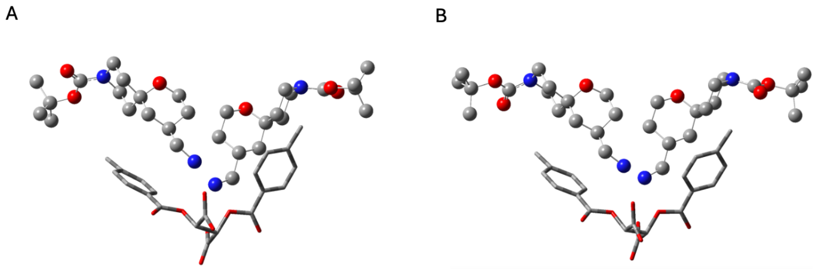

(A) Energy minimized conformers 1 and 3 (-489.99 kcalmol$^{-1}$). (B) Energy minimized conformers 2 and 4 (-485.86 kcalmol$^{-1}$). Amine **2** is shown in stick and ball representation. D-DTTA is shown as sticks only. The BSSE interaction energy difference between A and B (0.9 kcalmol$^{-1}$) arises from the C–N bond rotation of the Boc amide.

**Figure S5.** Enantioselectivity of **4**

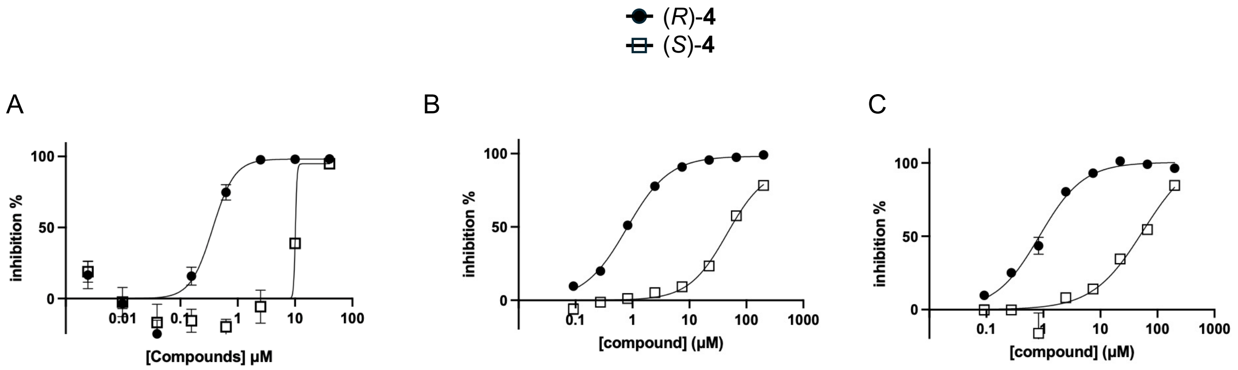

CHIKV-nLuc
(*R*)-**4** $EC_{50}$ = 0.36 µM
(*S*)-**4** $EC_{50}$ = 10 µM

ATPase
(*R*)-**4** $IC_{50}$ = 0.79 µM
(*S*)-**4** $IC_{50}$ = 46 µM

RNA Unwinding
(*R*)-**4** $IC_{50}$ = 0.85 µM
(*S*)-**4** $IC_{50}$ = 56 µM

**Figure S6.** ¹H NMR (500 MHz, $CD_3OD$) for ($R$-**2**)$_2$•D-DTTA

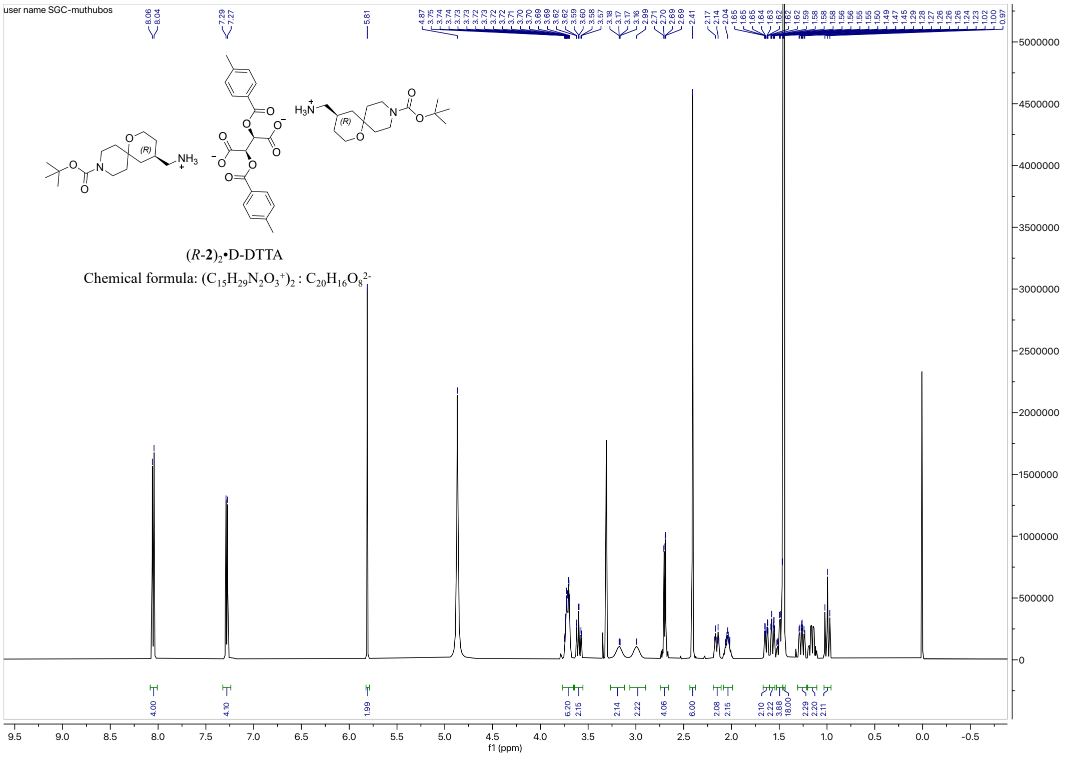

**Figure S7.** ¹³C NMR (126 MHz, $CD_3OD$) for ($R$-**2**)$_2$•D-DTTA

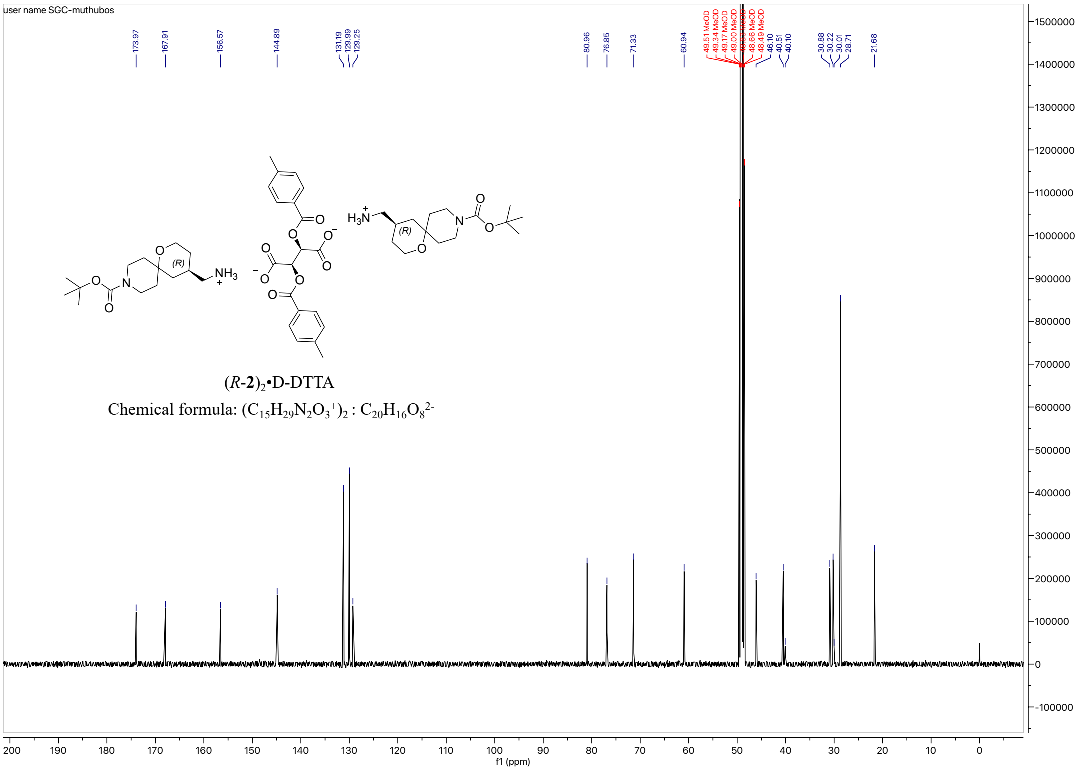

**Figure S8.** $^{1}$H NMR (400 MHz, DMSO-d$_6$) for (*R*)-**3**

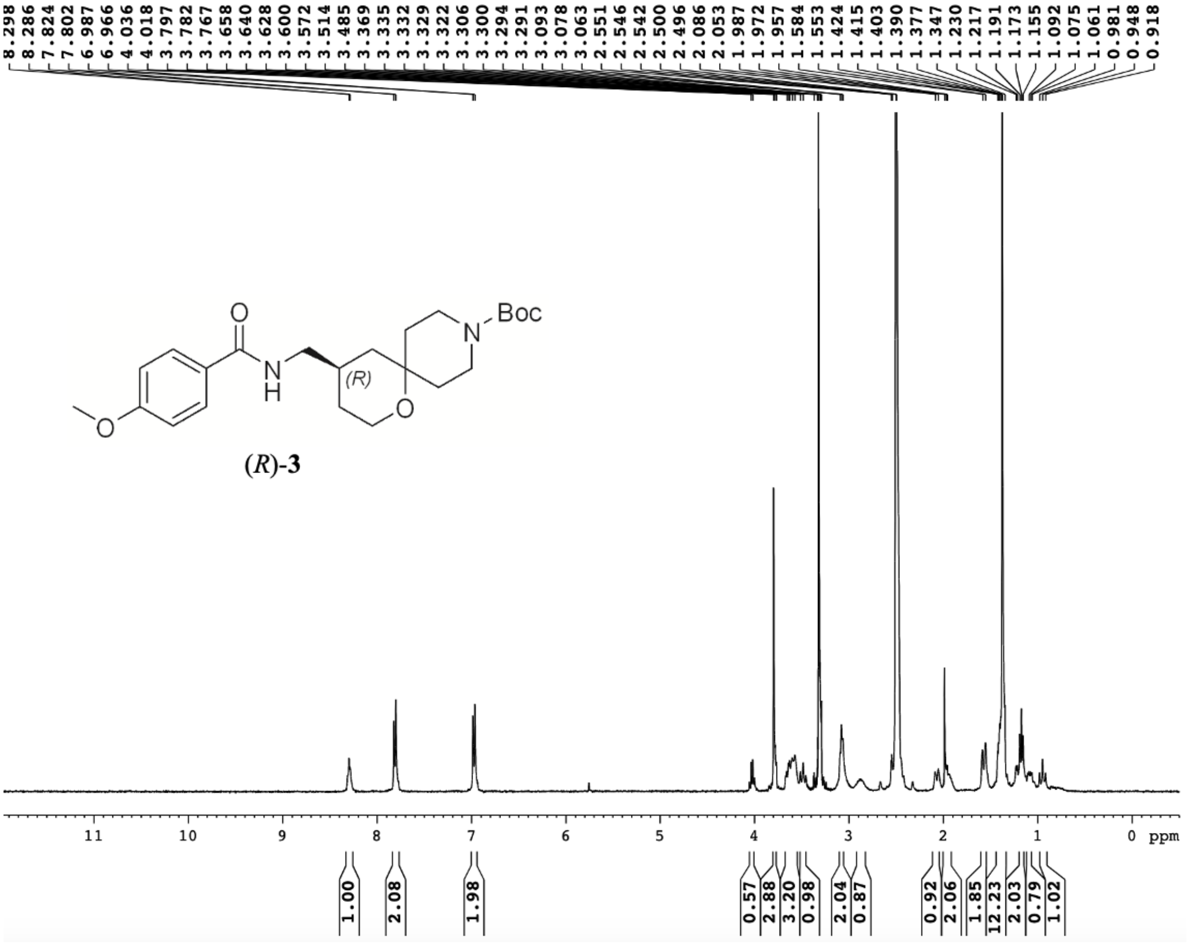

**Figure S9.** Chiral SFC trace of (*R*)-**3**

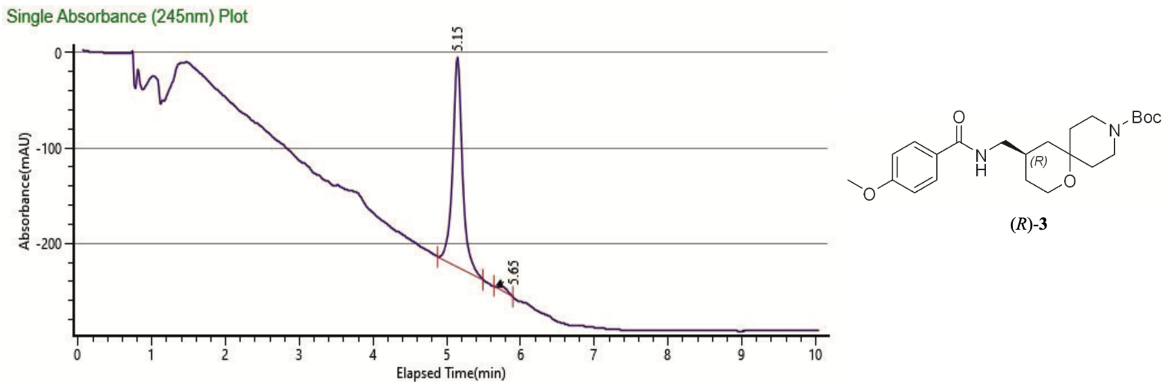

| Peak # | Ret. Time | Area | Height | Area % |
|---|---|---|---|---|
| 1 | 5.15 min | 2024.9507 | 220.0006 | 98.2549 |
| 2 | 5.65 min | 35.9657 | 0.0000 | 1.7451 |

**Figure S10.** Chiral SFC trace of (*S*)-**3**

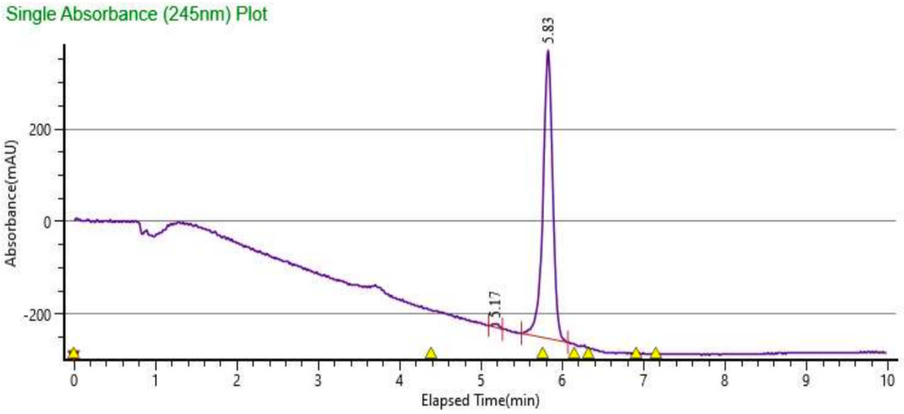

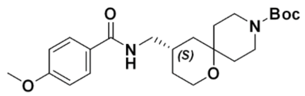

(*S*)-**3**

| Peak # | Ret. Time | Area | Height | Area % |
|---|---|---|---|---|
| 1 | 5.17 min | 41.9643 | 7.2548 | 0.8101 |
| 2 | 5.83 min | 5137.8781 | 622.3857 | 99.1899 |

**Figure S11.** $^{1}$H NMR (500 MHz, $CD_3CN$) for *R*-(+)-**1**

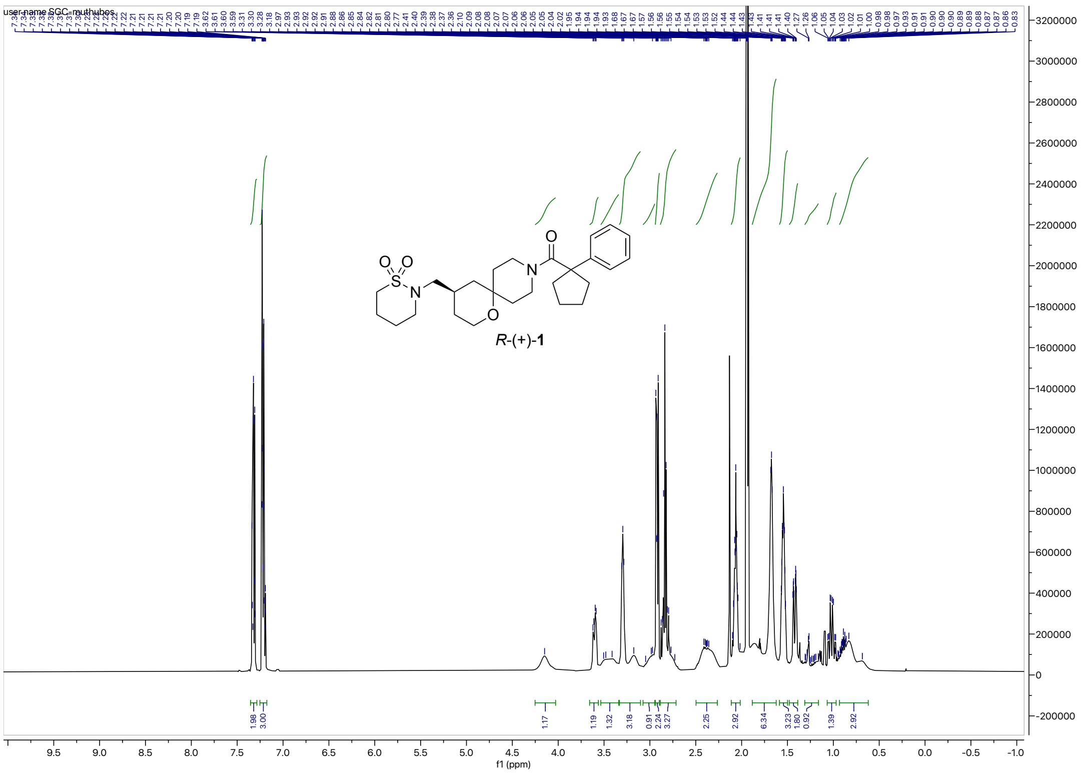

**Figure S12.** $^{13}$C NMR (126 MHz, $CD_3CN$) for *R*-(+)-**1**

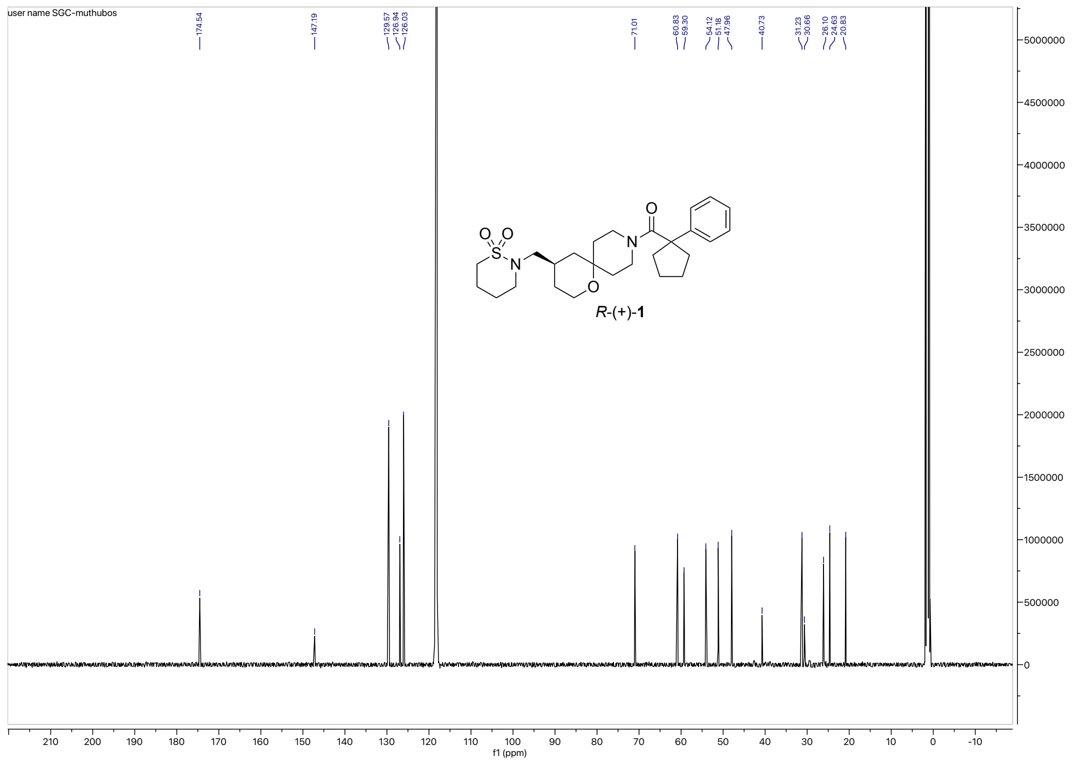

**Figure S13.** HPLC trace (220 nm) for *R*-(+)-**1**

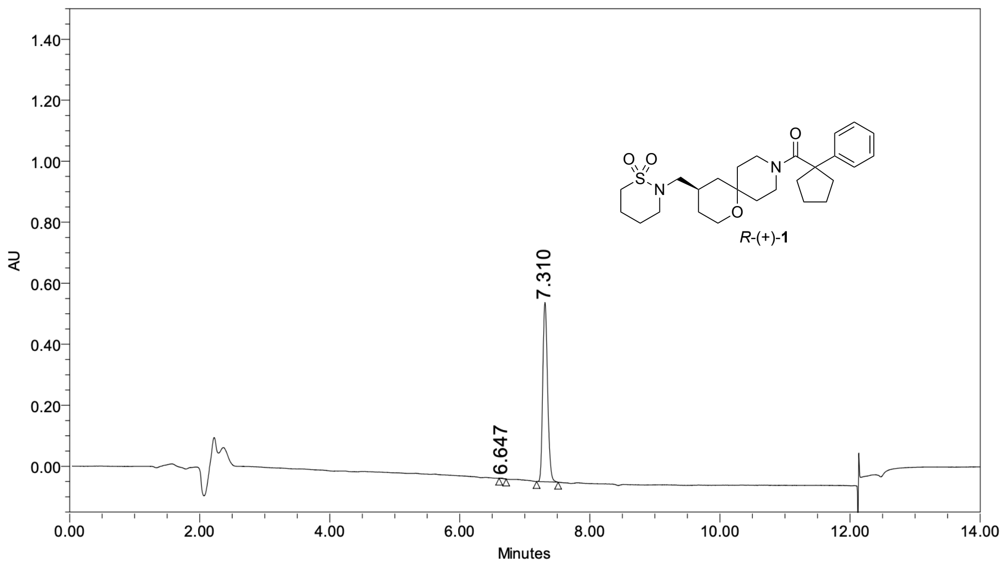

**Peak Results**

| | Name | RT | Height (μV) | Area (μV*sec) | % Area |
|---|---|---|---|---|---|
| 1 | | 6.647 | 1630 | 4924 | 0.16 |
| 2 | | 7.310 | 587636 | 3040581 | 99.84 |

**Figure S14.** Chiral HPLC (230 nm) of *R*-(+)-**1**

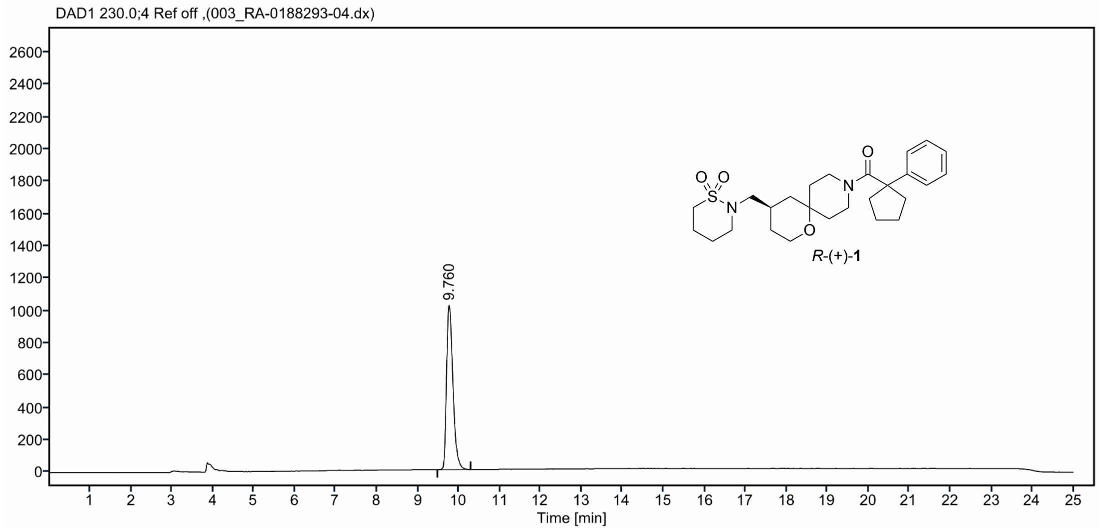

| Sr.No. | RT | Area | Height | % Area |
|---|---|---|---|---|
| 1 | 9.76 | 11005 | 1017 | 100.00 |

**Figure S15.** $^{1}H$ NMR (500 MHz, $CD_3CN$) for *S*-(-)-**1**

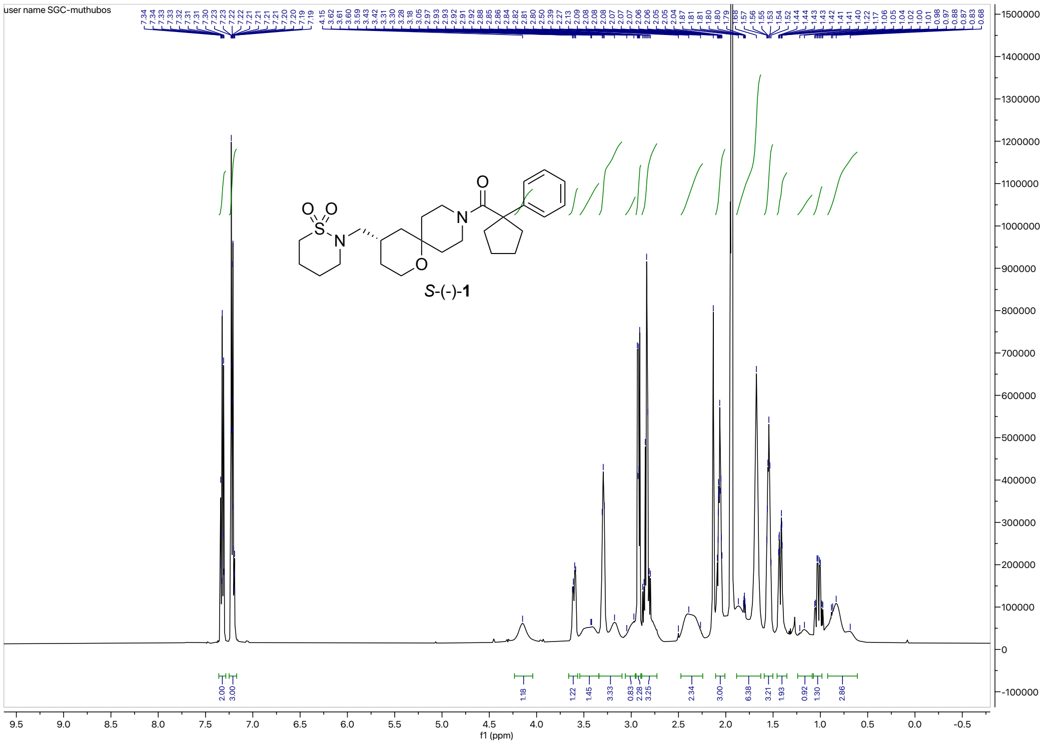

**Figure S16.** $^{13}C$ NMR (126 MHz, $CD_3CN$) for *S*-(-)-**1**

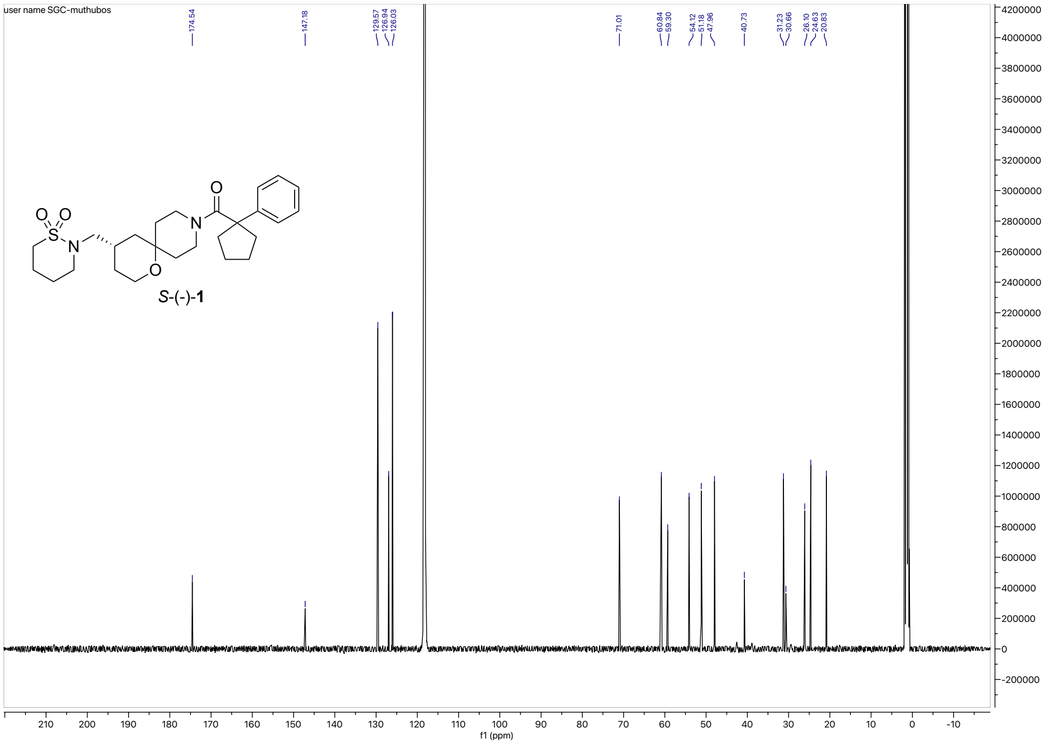

**Figure S17.** HPLC trace (UV- 220 nm) for *S*-(-)-**1**

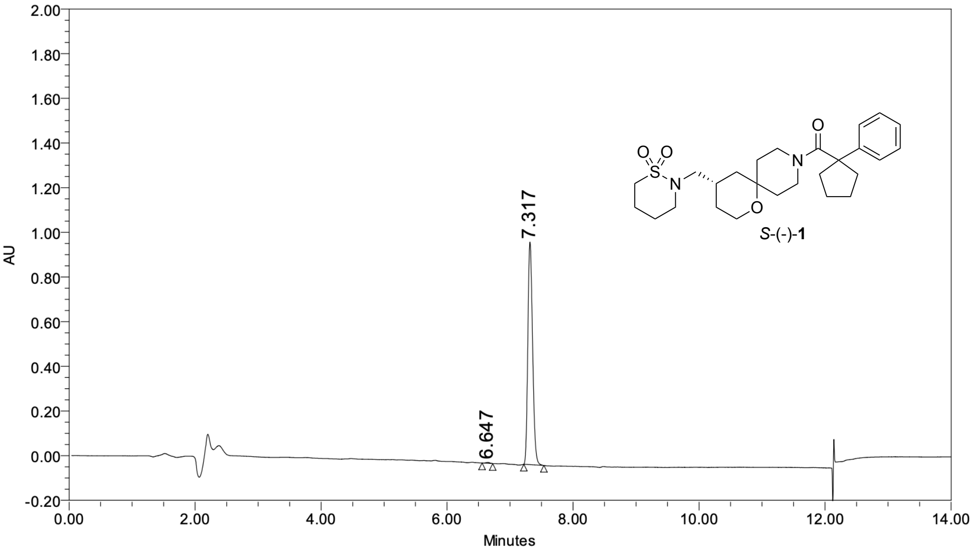

**Peak Results**

| | Name | RT | Height (µV) | Area (µV*sec) | % Area |
|---|---|---|---|---|---|
| 1 | | 6.647 | 4026 | 16308 | 0.31 |
| 2 | | 7.317 | 997795 | 5228748 | 99.69 |

**Figure S18.** Chiral HPLC (220 nm) of *S*-(-)-**1**

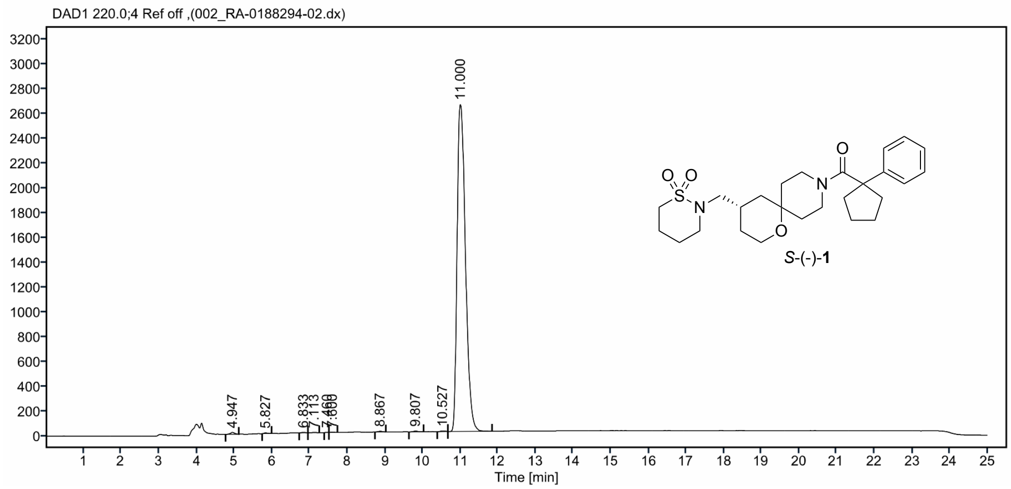

### Peak Results

| Sr. No | RT | % Area |
|---|---|---|
| 1 | 9.81 | 0.12 |
| 2 | 11.00 | 99.88 |

**Figure S19.** $^{1}$H NMR (500 MHz, DMSO-d$_6$) for (*R*)-**4**

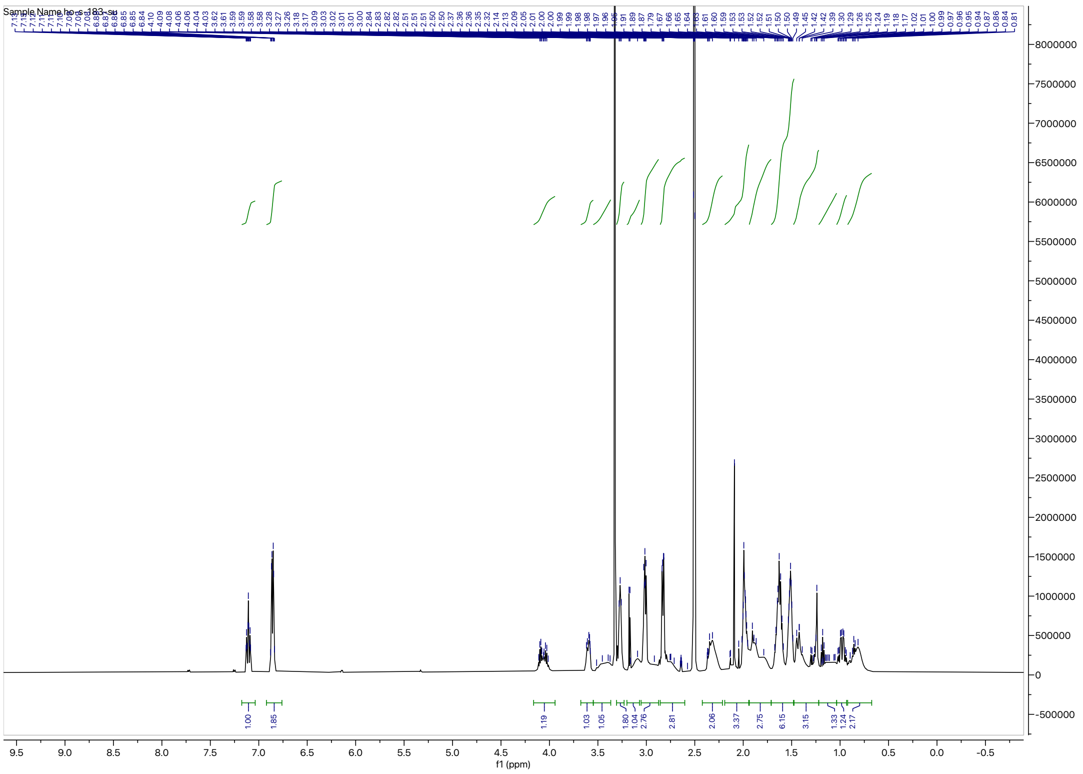

**Figure S20.** $^{13}$C NMR (126 MHz, DMSO-d$_6$) for (*R*)-**4**

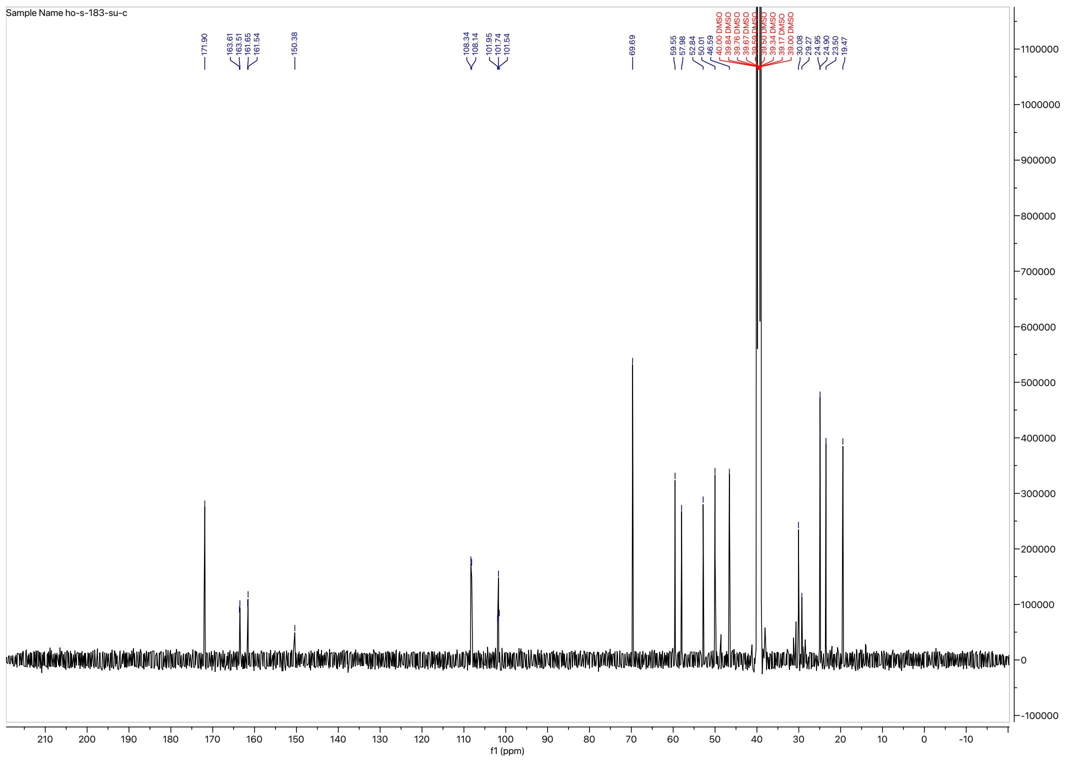

**Figure S21.** HRMS of (*R*)-**4**

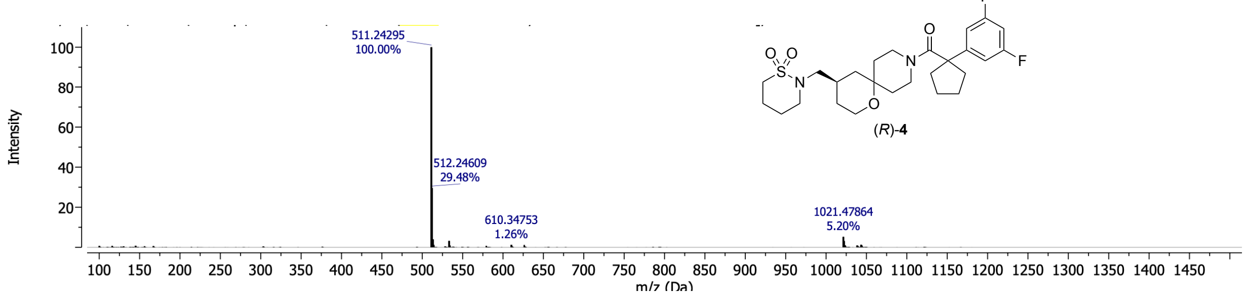

**Figure S22.** HPLC trace (254 nm) for (*R*)-**4**

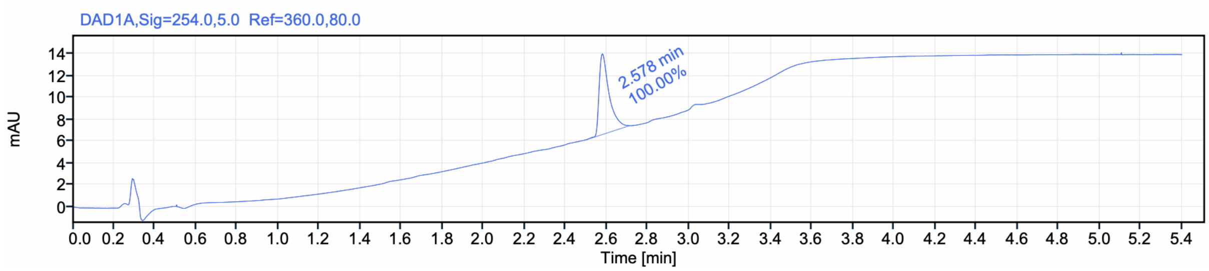

Signal Name DAD1A

| RT (min) | Signal description | Area | Area% | Peak Resolution USP |
|---|---|---|---|---|
| 2.578 | DAD1A,Sig=254.0,5.0 Ref=360.0,80.0 | 25.0 | 100.00 | |

**Figure S23.** HRMS of (*S*)-**4**

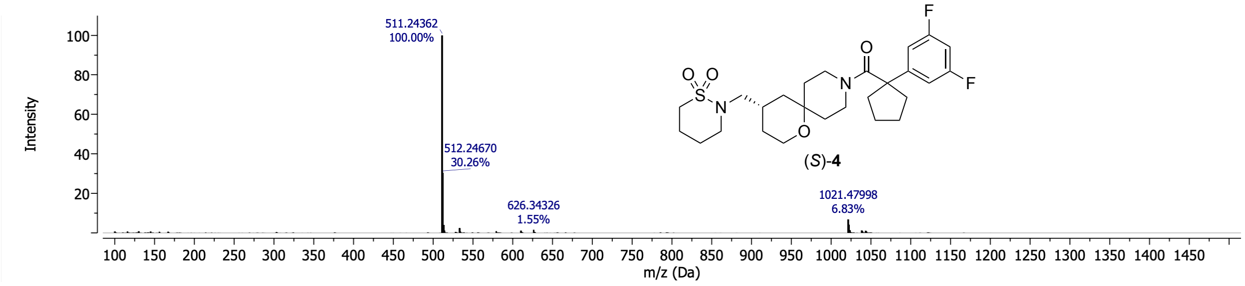

**Figure S24.** HPLC trace (UV- 254 nm) for (*S*)-**4**

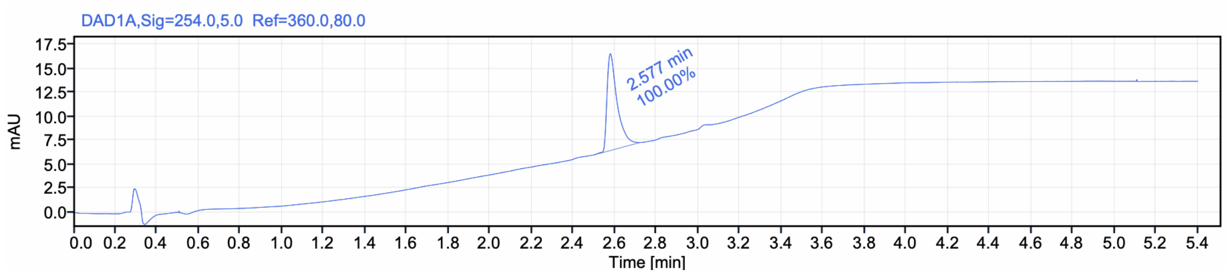

**Table S1.** Crystal data and structure refinement for (*R*-**2**)$_2$•D-DTTA

| | | |
|---|---|---|
| Empirical formula | C50 H74 N4 O14 | |
| Formula weight | 955.16 | |
| Crystal color, shape, size | colorless plate, 0.210 x 0.070 x 0.020 $mm^3$ | |
| Temperature | 150 K | |
| Wavelength | 1.54180 Å | |
| Crystal system, space group | Monoclinic, C 1 2 1 | |
| Unit cell dimensions | a = 16.8845(5) Å | $\alpha$= 90°. |
| | b = 6.9690(2) Å | $\beta$= 102.750(2)°. |
| | c = 25.3646(8) Å | $\gamma$ = 90°. |
| Volume | 2911.01(15) $Å^3$ | |
| Z | 2 | |
| Density (calculated) | 1.090 $Mg/m^3$ | |
| Absorption coefficient | 0.652 $mm^{-1}$ | |
| F(000) | 1028.002 | |
| Data collection | | |
| Diffractometer | Bruker D8 VENTURE, Bruker | |
| Theta range for data collection | 1.786 to 68.395°. | |
| Index ranges | -20<=h<=19, -8<=k<=8, -30<=l<=30 | |
| Reflections collected | 27729 | |
| Independent reflections | 5327 [R(int) = 0.058] | |
| Observed Reflections | 4940 | |
| Completeness to theta = 68.395° | 99.7 % | |
| Solution and Refinement | | |
| Absorption correction | Semi-empirical from equivalents | |
| Max. and min. transmission | 0.99 and 0.96 | |
| Solution | Intrinsic phasing methods | |
| Refinement method | Full-matrix least-squares on $F^2$ | |
| Weighting scheme | w = $[\sigma^2 Fo^2 + AP^2 + BP]^{-1}$, with | |
| | P = $(Fo^2 + 2\ Fc^2)/3$, A = 0.222, B = 2.098 | |
| Data / restraints / parameters | 5327 / 602 / 505 | |
| Goodness-of-fit on $F^2$ | 0.9903 | |
| Final R indices [I>2sigma(I)] | R1 = 0.0865, wR2 = 0.2515 | |
| R indices (all data) | R1 = 0.0930, wR2 = 0.2602 | |
| Absolute structure parameter | 0.08(6) | |
| Largest diff. peak and hole | 0.90 and -0.44 e.$Å^{-3}$ | |

**Table S2.** Atomic coordinates for ($R$-**2**)$_2$•D-DTTA

| | x | y | z | U(eq) |
|---|---|---|---|---|
| O1 | 8646(2) | 9356(8) | 6949(1) | 53 |
| O2 | 7712(5) | 11154(10) | 8811(3) | 128 |
| O3 | 8980(5) | 9802(12) | 9122(3) | 118 |
| O4 | 5296(1) | 1782(8) | 5539(1) | 33 |
| O5 | 6649(1) | 1241(8) | 5117(1) | 38 |
| O6 | 6276(1) | -1762(8) | 4874(1) | 37 |
| O7 | 6125(1) | 397(8) | 6255(1) | 42 |
| O202 | 7875(6) | 12826(14) | 8590(3) | 127 |
| O203 | 9157(6) | 11687(14) | 8965(4) | 134 |
| N1 | 8501(4) | 10449(9) | 8168(2) | 91 |
| N2 | 7104(2) | 5002(8) | 5383(1) | 35 |
| N201 | 8307(4) | 10016(11) | 8222(2) | 80 |
| C1 | 8183(2) | 9874(9) | 6434(2) | 46 |
| C2 | 8039(2) | 8152(9) | 6056(2) | 42 |
| C3 | 7589(2) | 6577(8) | 6293(1) | 38 |
| C4 | 8056(2) | 6156(9) | 6870(2) | 50 |
| C5 | 8260(2) | 7953(8) | 7225(1) | 55 |
| C6 | 7541(3) | 8837(17) | 7411(4) | 62 |
| C7 | 7777(6) | 10619(15) | 7758(2) | 69 |
| C8 | 9166(4) | 9370(17) | 8067(3) | 86 |
| C9 | 8939(6) | 7580(13) | 7722(3) | 66 |
| C10 | 7504(2) | 4738(8) | 5972(1) | 38 |
| C11 | 8390(4) | 10398(10) | 8727(2) | 114 |
| C12 | 6982(5) | 10035(18) | 8710(5) | 134 |
| C13 | 6261(7) | 11210(30) | 8390(10) | 168 |
| C14 | 6778(13) | 9390(40) | 9245(8) | 167 |
| C15 | 7079(9) | 8230(30) | 8379(11) | 173 |
| C16 | 5358(2) | 53(8) | 5244(1) | 30 |
| C17 | 6165(2) | -151(8) | 5062(1) | 31 |
| C18 | 5711(2) | 1749(8) | 6059(1) | 32 |
| C19 | 5599(2) | 3516(9) | 6357(1) | 36 |
| C20 | 5171(2) | 5091(8) | 6105(1) | 36 |
| C21 | 5073(2) | 6699(9) | 6417(2) | 42 |
| C22 | 5382(2) | 6756(9) | 6961(2) | 47 |
| C23 | 5819(3) | 5173(9) | 7210(2) | 54 |
| C24 | 5930(2) | 3552(9) | 6907(2) | 45 |
| C25 | 5244(3) | 8465(10) | 7293(2) | 60 |
| C112 | 8790(9) | 8900(30) | 9587(6) | 139 |
| C113 | 9220(20) | 6950(40) | 9695(17) | 156 |

| | | | | |
|---|---|---|---|---|
| C114 | 7869(10) | 8560(60) | 9500(10) | 155 |
| C115 | 9060(20) | 10170(70) | 10092(5) | 144 |
| C206 | 7491(3) | 8742(17) | 7365(3) | 57 |
| C207 | 7663(6) | 10354(16) | 7770(3) | 85 |
| C208 | 9006(4) | 9014(16) | 8155(3) | 83 |
| C209 | 8850(6) | 7390(12) | 7749(3) | 70 |
| C211 | 8446(4) | 11521(12) | 8631(3) | 117 |
| C212 | 7802(8) | 13910(17) | 9057(4) | 139 |
| C213 | 7733(17) | 12560(30) | 9527(6) | 168 |
| C214 | 7038(13) | 15190(30) | 8928(7) | 165 |
| C215 | 8548(13) | 15210(30) | 9245(9) | 172 |
| C312 | 9790(8) | 12760(30) | 8806(8) | 141 |
| C313 | 9830(30) | 14810(40) | 9037(18) | 143 |
| C314 | 10620(7) | 11790(80) | 9010(20) | 156 |
| C315 | 9635(18) | 12900(70) | 8185(9) | 144 |

Atomic coordinates for (*R*-**2**)$_2$•D-DTTA (x $10^4$) and equivalent isotropic displacement parameters ($Å^2$x $10^3$) for (*R*-**2**)$_2$•D-DTTA. U(eq) is defined as one third of the trace of the orthogonalized $U^{ij}$ tensor.

**Table S3.** Bond lengths [Å] and angles [°] for (*R*-**2**)$_2$•D-DTTA.

| | | | |
|---|---|---|---|
| O1-C1 | 1.412(5) | O1-C5 | 1.441(5) |
| O2-C11 | 1.320(5) | O2-C12 | 1.433(4) |
| O3-C11 | 1.315(5) | O3-C112 | 1.433(4) |
| O4-C16 | 1.435(3) | O4-C18 | 1.350(4) |
| O5-C17 | 1.256(4) | O6-C17 | 1.250(4) |
| O7-C18 | 1.212(4) | O202-C211 | 1.313(5) |
| O202-C212 | 1.432(4) | O203-C211 | 1.313(5) |
| O203-C312 | 1.434(4) | N1-C7 | 1.425(4) |
| N1-C8 | 1.422(4) | N1-C11 | 1.471(5) |
| N2-C10 | 1.509(4) | N2-H1 | 0.893(19) |
| N2-H2 | 0.888(19) | N2-H3 | 0.904(19) |
| N201-C207 | 1.416(5) | N201-C208 | 1.415(4) |
| N201-C211 | 1.457(4) | C1-C2 | 1.521(5) |
| C1-H11 | 0.950 | C1-H12 | 0.950 |
| C2-C3 | 1.531(5) | C2-H21 | 0.950 |
| C2-H22 | 0.950 | C3-C4 | 1.529(5) |
| C3-C10 | 1.508(5) | C3-H31 | 0.950 |
| C4-C5 | 1.537(5) | C4-H41 | 0.950 |
| C4-H42 | 0.950 | C5-C6 | 1.526(6) |
| C5-C9 | 1.526(6) | C5-C206 | 1.521(5) |
| C5-C209 | 1.525(5) | C6-C7 | 1.523(6) |
| C6-H61 | 0.950 | C6-H62 | 0.950 |
| C7-H71 | 0.950 | C7-H73 | 0.950 |
| C8-C9 | 1.524(6) | C8-H81 | 0.950 |
| C8-H82 | 0.950 | C9-H91 | 0.950 |
| C9-H92 | 0.950 | C10-H101 | 0.950 |
| C10-H102 | 0.950 | C12-C13 | 1.543(5) |
| C12-C14 | 1.539(5) | C12-C15 | 1.540(5) |
| C13-H131 | 0.950 | C13-H132 | 0.950 |
| C13-H133 | 0.950 | C14-H141 | 0.950 |
| C14-H142 | 0.950 | C14-H143 | 0.950 |
| C15-H151 | 0.950 | C15-H152 | 0.950 |
| C15-H153 | 0.950 | C16-C16#1 | 1.526(5) |
| C16-C17 | 1.538(4) | C16-H161 | 0.950 |
| C18-C19 | 1.478(4) | C19-C20 | 1.390(5) |
| C19-C24 | 1.382(5) | C20-C21 | 1.403(5) |
| C20-H201 | 0.950 | C21-C22 | 1.362(6) |
| C21-H211 | 0.950 | C22-C23 | 1.398(6) |
| C22-C25 | 1.506(5) | C23-C24 | 1.402(6) |
| C23-H231 | 0.950 | C24-H241 | 0.950 |
| C25-H251 | 0.950 | C25-H252 | 0.950 |

| | | | |
|---|---|---|---|
| C25-H253 | 0.950 | C112-C113 | 1.540(5) |
| C112-C114 | 1.541(5) | C112-C115 | 1.541(5) |
| C113-H1131 | 0.950 | C113-H1132 | 0.950 |
| C113-H1133 | 0.950 | C114-H1141 | 0.950 |
| C114-H1142 | 0.950 | C114-H1143 | 0.950 |
| C115-H1151 | 0.950 | C115-H1152 | 0.950 |
| C115-H1153 | 0.950 | C206-C207 | 1.507(6) |
| C206-H2061 | 0.950 | C206-H2062 | 0.950 |
| C207-H2071 | 0.950 | C207-H2073 | 0.950 |
| C208-C209 | 1.514(6) | C208-H2081 | 0.950 |
| C208-H2082 | 0.950 | C209-H2091 | 0.950 |
| C209-H2092 | 0.950 | C212-C213 | 1.543(5) |
| C212-C214 | 1.543(5) | C212-C215 | 1.540(5) |
| C213-H2131 | 0.950 | C213-H2132 | 0.950 |
| C213-H2133 | 0.950 | C214-H2141 | 0.950 |
| C214-H2142 | 0.950 | C214-H2143 | 0.950 |
| C215-H2151 | 0.950 | C215-H2152 | 0.950 |
| C215-H2153 | 0.950 | C312-C313 | 1.541(5) |
| C312-C314 | 1.541(5) | C312-C315 | 1.541(5) |
| C313-H3131 | 0.950 | C313-H3132 | 0.950 |
| C313-H3133 | 0.950 | C314-H3141 | 0.950 |
| C314-H3142 | 0.950 | C314-H3143 | 0.950 |
| C315-H3151 | 0.950 | C315-H3152 | 0.950 |
| C315-H3153 | 0.950 | | |

| | | | |
|---|---|---|---|
| C1-O1-C5 | 114.0(3) | C11-O2-C12 | 119.73(7) |
| C11-O3-C112 | 119.73(7) | C16-O4-C18 | 114.5(2) |
| C211-O202-C212 | 119.77(7) | C211-O203-C312 | 119.73(7) |
| C7-N1-C8 | 119.87(5) | C7-N1-C11 | 115.74(7) |
| C8-N1-C11 | 115.73(7) | C10-N2-H1 | 107.0(16) |
| C10-N2-H2 | 109.3(16) | H1-N2-H2 | 109.0(17) |
| C10-N2-H3 | 112.1(16) | H1-N2-H3 | 110.2(17) |
| H2-N2-H3 | 109.2(17) | C207-N201-C208 | 119.82(5) |
| C207-N201-C211 | 115.60(7) | C208-N201-C211 | 115.63(7) |
| O1-C1-C2 | 111.0(3) | O1-C1-H11 | 109.0 |
| C2-C1-H11 | 109.1 | O1-C1-H12 | 109.1 |
| C2-C1-H12 | 109.1 | H11-C1-H12 | 109.5 |
| C1-C2-C3 | 110.0(3) | C1-C2-H21 | 109.4 |
| C3-C2-H21 | 109.3 | C1-C2-H22 | 109.2 |
| C3-C2-H22 | 109.4 | H21-C2-H22 | 109.5 |
| C2-C3-C4 | 108.4(3) | C2-C3-C10 | 113.2(3) |
| C4-C3-C10 | 108.7(3) | C2-C3-H31 | 108.8 |
| C4-C3-H31 | 108.9 | C10-C3-H31 | 108.8 |

| | | | |
|---|---|---|---|
| C3-C4-C5 | 114.0(3) | C3-C4-H41 | 108.4 |
| C5-C4-H41 | 108.4 | C3-C4-H42 | 108.3 |
| C5-C4-H42 | 108.2 | H41-C4-H42 | 109.5 |
| C4-C5-O1 | 109.5(3) | C4-C5-C6 | 114.8(4) |
| O1-C5-C6 | 110.6(5) | C4-C5-C9 | 112.1(3) |
| O1-C5-C9 | 100.3(5) | C6-C5-C9 | 108.53(7) |
| C4-C5-O1 | 109.5(3) | C4-C5-C206 | 109.8(4) |
| O1-C5-C206 | 111.6(5) | C4-C5-C209 | 108.7(3) |
| O1-C5-C209 | 108.6(5) | C206-C5-C209 | 108.56(7) |
| C5-C6-C7 | 112.52(7) | C5-C6-H61 | 108.4 |
| C7-C6-H61 | 108.6 | C5-C6-H62 | 108.9 |
| C7-C6-H62 | 108.9 | H61-C6-H62 | 109.5 |
| C6-C7-H71 | 109.4 | C6-C7-H73 | 109.3 |
| H71-C7-H73 | 109.5 | C9-C8-H81 | 109.4 |
| C9-C8-H82 | 109.3 | H81-C8-H82 | 109.5 |
| C5-C9-C8 | 112.53(7) | C5-C9-H91 | 108.9 |
| C8-C9-H91 | 108.9 | C5-C9-H92 | 108.4 |
| C8-C9-H92 | 108.6 | H91-C9-H92 | 109.5 |
| N2-C10-C3 | 113.6(3) | N2-C10-H101 | 108.5 |
| C3-C10-H101 | 108.4 | N2-C10-H102 | 108.3 |
| C3-C10-H102 | 108.6 | H101-C10-H102 | 109.5 |
| O2-C11-O3 | 122.77(6) | O2-C12-C13 | 110.62(5) |
| O2-C12-C14 | 110.61(5) | C13-C12-C14 | 108.30(5) |
| O2-C12-C15 | 110.61(5) | C13-C12-C15 | 108.31(5) |
| C14-C12-C15 | 108.30(5) | C12-C13-H131 | 110.0 |
| C12-C13-H132 | 109.0 | H131-C13-H132 | 109.5 |
| C12-C13-H133 | 109.4 | H131-C13-H133 | 109.5 |
| H132-C13-H133 | 109.5 | C12-C14-H141 | 108.8 |
| C12-C14-H142 | 110.1 | H141-C14-H142 | 109.5 |
| C12-C14-H143 | 109.4 | H141-C14-H143 | 109.5 |
| H142-C14-H143 | 109.5 | C12-C15-H151 | 109.8 |
| C12-C15-H152 | 109.9 | H151-C15-H152 | 109.5 |
| C12-C15-H153 | 108.7 | H151-C15-H153 | 109.5 |
| H152-C15-H153 | 109.5 | C16#1-C16-O4 | 106.69(18) |
| C16#1-C16-C17 | 110.7(3) | O4-C16-C17 | 113.9(2) |
| C16#1-C16-H161 | 108.5 | O4-C16-H161 | 108.4 |
| C17-C16-H161 | 108.5 | C16-C17-O5 | 119.5(3) |
| C16-C17-O6 | 114.5(3) | O5-C17-O6 | 126.0(3) |
| O4-C18-O7 | 122.7(3) | O4-C18-C19 | 112.9(3) |
| O7-C18-C19 | 124.4(3) | C18-C19-C20 | 122.3(3) |
| C18-C19-C24 | 117.5(3) | C20-C19-C24 | 120.2(3) |
| C19-C20-C21 | 119.0(3) | C19-C20-H201 | 120.5 |
| C21-C20-H201 | 120.5 | C20-C21-C22 | 121.9(3) |

| | | | |
|---|---|---|---|
| C20-C21-H211 | 119.1 | C22-C21-H211 | 119.0 |
| C21-C22-C23 | 118.7(4) | C21-C22-C25 | 121.1(4) |
| C23-C22-C25 | 120.2(4) | C22-C23-C24 | 120.7(3) |
| C22-C23-H231 | 119.5 | C24-C23-H231 | 119.8 |
| C23-C24-C19 | 119.6(3) | C23-C24-H241 | 120.1 |
| C19-C24-H241 | 120.3 | C22-C25-H251 | 109.5 |
| C22-C25-H252 | 109.6 | H251-C25-H252 | 109.5 |
| C22-C25-H253 | 109.3 | H251-C25-H253 | 109.5 |
| H252-C25-H253 | 109.5 | O3-C112-C113 | 110.61(5) |
| O3-C112-C114 | 110.61(5) | C113-C112-C114 | 108.31(5) |
| O3-C112-C115 | 110.61(5) | C113-C112-C115 | 108.31(5) |
| C114-C112-C115 | 108.31(5) | C112-C113-H1131 | 109.7 |
| C112-C113-H1132 | 107.6 | H1131-C113-H1132 | 109.5 |
| C112-C113-H1133 | 111.0 | H1131-C113-H1133 | 109.5 |
| H1132-C113-H1133 | 109.5 | C112-C114-H1141 | 109.5 |
| C112-C114-H1142 | 111.2 | H1141-C114-H1142 | 109.5 |
| C112-C114-H1143 | 107.7 | H1141-C114-H1143 | 109.5 |
| H1142-C114-H1143 | 109.5 | C112-C115-H1151 | 109.4 |
| C112-C115-H1152 | 111.2 | H1151-C115-H1152 | 109.5 |
| C112-C115-H1153 | 107.8 | H1151-C115-H1153 | 109.5 |
| H1152-C115-H1153 | 109.5 | C5-C206-C207 | 112.51(7) |
| C5-C206-H2061 | 109.0 | C207-C206-H2061 | 108.4 |
| C5-C206-H2062 | 108.5 | C207-C206-H2062 | 108.9 |
| H2061-C206-H2062 | 109.5 | C206-C207-H2071 | 109.6 |
| C206-C207-H2073 | 109.2 | H2071-C207-H2073 | 109.5 |
| C209-C208-H2081 | 109.4 | C209-C208-H2082 | 109.1 |
| H2081-C208-H2082 | 109.5 | C5-C209-C208 | 112.51(7) |
| C5-C209-H2091 | 109.3 | C208-C209-H2091 | 109.1 |
| C5-C209-H2092 | 108.0 | C208-C209-H2092 | 108.4 |
| H2091-C209-H2092 | 109.5 | O203-C211-O202 | 122.74(6) |
| O202-C212-C213 | 110.62(5) | O202-C212-C214 | 110.60(5) |
| C213-C212-C214 | 108.31(5) | O202-C212-C215 | 110.61(5) |
| C213-C212-C215 | 108.31(5) | C214-C212-C215 | 108.31(5) |
| C212-C213-H2131 | 110.9 | C212-C213-H2132 | 112.0 |
| H2131-C213-H2132 | 109.5 | C212-C213-H2133 | 105.4 |
| H2131-C213-H2133 | 109.5 | H2132-C213-H2133 | 109.5 |
| C212-C214-H2141 | 110.0 | C212-C214-H2142 | 106.9 |
| H2141-C214-H2142 | 109.5 | C212-C214-H2143 | 111.5 |
| H2141-C214-H2143 | 109.5 | H2142-C214-H2143 | 109.5 |
| C212-C215-H2151 | 107.7 | C212-C215-H2152 | 107.2 |
| H2151-C215-H2152 | 109.5 | C212-C215-H2153 | 113.5 |
| H2151-C215-H2153 | 109.5 | H2152-C215-H2153 | 109.5 |
| O203-C312-C313 | 110.61(5) | O203-C312-C314 | 110.61(5) |

| | | | |
|---|---|---|---|
| C313-C312-C314 | 108.31(5) | O203-C312-C315 | 110.61(5) |
| C313-C312-C315 | 108.31(5) | C314-C312-C315 | 108.31(5) |
| C312-C313-H3131 | 107.7 | C312-C313-H3132 | 111.6 |
| H3131-C313-H3132 | 109.5 | C312-C313-H3133 | 109.1 |
| H3131-C313-H3133 | 109.5 | H3132-C313-H3133 | 109.5 |
| C312-C314-H3141 | 111.7 | C312-C314-H3142 | 107.0 |
| H3141-C314-H3142 | 109.5 | C312-C314-H3143 | 109.7 |
| H3141-C314-H3143 | 109.5 | H3142-C314-H3143 | 109.5 |
| C312-C315-H3151 | 109.6 | C312-C315-H3152 | 112.0 |
| H3151-C315-H3152 | 109.5 | C312-C315-H3153 | 106.7 |
| H3151-C315-H3153 | 109.5 | H3152-C315-H3153 | 109.5 |

Symmetry transformations used to generate equivalent atoms: #1 -x+1,y,-z+1

**Table S4.** Anisotropic displacement parameters for ($R$-**2**)$_2$•D-DTTA.

| — | $U^{11}$ | $U^{22}$ | $U^{33}$ | $U^{23}$ | $U^{13}$ | $U^{12}$ |
|---|---|---|---|---|---|---|
| — | | | | | | |
| O1 | 37(1) | 47(2) | 69(2) | -8(1) | -3(1) | -1(1) |
| O2 | 134(6) | 145(6) | 102(5) | -22(5) | 20(5) | 3(6) |
| O3 | 131(5) | 131(5) | 78(4) | -22(4) | -8(4) | 7(5) |
| O4 | 29(1) | 28(1) | 42(1) | -2(1) | 6(1) | 4(1) |
| O5 | 30(1) | 28(1) | 60(1) | 0(1) | 14(1) | -4(1) |
| O6 | 29(1) | 27(1) | 56(1) | -3(1) | 9(1) | 2(1) |
| O7 | 37(1) | 36(1) | 49(1) | 1(1) | 1(1) | 4(1) |
| O202 | 136(6) | 137(6) | 103(5) | -16(5) | 17(5) | 12(6) |
| O203 | 132(6) | 157(7) | 102(5) | -16(5) | 0(5) | 4(6) |
| N1 | 92(3) | 98(4) | 75(3) | -18(3) | 2(3) | 0(4) |
| N2 | 28(1) | 26(1) | 53(1) | -2(1) | 12(1) | 1(1) |
| N201 | 81(3) | 87(4) | 64(3) | -17(3) | -3(3) | 5(3) |
| C1 | 35(2) | 34(2) | 66(2) | -1(2) | 7(1) | -2(1) |
| C2 | 32(2) | 33(2) | 62(2) | 5(1) | 10(1) | 0(1) |
| C3 | 31(1) | 32(2) | 50(2) | 4(1) | 10(1) | 1(1) |
| C4 | 43(2) | 41(2) | 63(2) | 6(2) | 6(2) | 9(2) |
| C5 | 52(2) | 53(2) | 53(2) | 3(2) | -3(2) | 10(2) |
| C6 | 60(4) | 71(4) | 54(4) | -7(4) | 11(4) | 5(4) |
| C7 | 72(4) | 77(5) | 60(4) | -23(4) | 17(4) | 3(4) |
| C8 | 85(3) | 95(4) | 71(3) | -14(3) | -1(3) | -1(4) |
| C9 | 56(4) | 67(4) | 61(4) | 2(4) | -13(4) | 6(4) |
| C10 | 31(1) | 31(2) | 53(2) | 4(1) | 10(1) | 0(1) |
| C11 | 126(7) | 110(7) | 87(6) | -25(5) | -13(6) | 1(6) |
| C12 | 140(6) | 150(6) | 104(5) | -16(6) | 11(5) | 6(6) |
| C13 | 170(10) | 181(10) | 139(9) | -9(9) | 3(9) | -10(10) |
| C14 | 177(10) | 178(10) | 141(9) | -3(9) | 26(9) | 15(10) |
| C15 | 175(10) | 190(10) | 143(10) | -7(10) | 13(9) | 5(10) |
| C16 | 27(1) | 23(1) | 41(2) | -1(1) | 7(1) | 1(1) |
| C17 | 25(1) | 26(1) | 42(1) | 2(1) | 6(1) | 2(1) |
| C18 | 24(1) | 33(2) | 38(1) | -1(1) | 6(1) | -1(1) |
| C19 | 28(1) | 35(2) | 45(2) | -5(1) | 8(1) | -4(1) |
| C20 | 30(1) | 35(2) | 44(2) | -2(1) | 9(1) | 0(1) |
| C21 | 32(1) | 34(2) | 61(2) | -3(2) | 13(1) | 0(1) |
| C22 | 45(2) | 40(2) | 58(2) | -10(2) | 14(2) | -3(2) |
| C23 | 53(2) | 54(2) | 51(2) | -9(2) | 4(2) | 4(2) |
| C24 | 43(2) | 42(2) | 48(2) | -2(2) | 3(1) | 4(2) |
| C25 | 64(2) | 51(2) | 67(2) | -18(2) | 18(2) | 2(2) |

| | | | | | | |
|---|---|---|---|---|---|---|
| C112 | 146(6) | 153(6) | 106(5) | -14(5) | 6(5) | 6(6) |
| C113 | 163(8) | 172(9) | 124(8) | -9(8) | 10(8) | 7(8) |
| C114 | 160(10) | 168(10) | 125(9) | -12(9) | 4(9) | -3(9) |
| C115 | 151(11) | 157(12) | 114(11) | -10(11) | 11(11) | 6(11) |
| C206 | 53(4) | 64(4) | 49(4) | -1(3) | 2(3) | 12(4) |
| C207 | 83(5) | 91(5) | 74(5) | -24(4) | 6(4) | 13(5) |
| C208 | 79(3) | 90(4) | 67(3) | -13(3) | -10(3) | 6(3) |
| C209 | 66(5) | 69(4) | 62(4) | -2(4) | -13(4) | 13(4) |
| C211 | 120(8) | 111(8) | 96(7) | -10(7) | -29(6) | 9(7) |
| C212 | 149(6) | 152(6) | 108(5) | -17(6) | 13(5) | 9(6) |
| C213 | 180(10) | 179(10) | 140(9) | -3(9) | 25(8) | 16(9) |
| C214 | 170(8) | 180(8) | 138(7) | -11(7) | 17(7) | 5(8) |
| C215 | 175(10) | 190(10) | 142(9) | -15(10) | 16(9) | 2(10) |
| C312 | 145(6) | 160(7) | 109(6) | -14(6) | 6(6) | 6(6) |
| C313 | 149(12) | 162(12) | 107(11) | -8(12) | 5(11) | 10(12) |
| C314 | 163(8) | 170(9) | 126(8) | -8(8) | 13(8) | 10(8) |
| C315 | 150(12) | 162(12) | 109(11) | -16(12) | 8(11) | 5(12) |

The anisotropic displacement factor exponent takes the form: $-2\pi^2[h^2 a^{*2}U^{11} + 2 h k a^* b^* U^{12}]$

**Table S5.** Hydrogen coordinates for (*R*-**2**)$_2$•D-DTTA

| | x | y | z | U(eq) |
|---|---|---|---|---|
| H1 | 6724(17) | 5910(40) | 5366(10) | 52(2) |
| H2 | 6866(19) | 3910(30) | 5253(10) | 53(2) |
| H3 | 7463(14) | 5350(50) | 5184(9) | 53(2) |
| H11 | 8466 | 10834 | 6284 | 54 |
| H12 | 7674 | 10368 | 6473 | 54 |
| H21 | 8546 | 7667 | 6012 | 51 |
| H22 | 7722 | 8539 | 5715 | 51 |
| H31 | 7062 | 7031 | 6305 | 45 |
| H41 | 8550 | 5533 | 6852 | 60 |
| H42 | 7736 | 5327 | 7035 | 60 |
| H61 | 7330 | 7909 | 7618 | 73 |
| H62 | 7134 | 9175 | 7102 | 73 |
| H71 | 7312 | 11112 | 7865 | 83 |
| H73 | 7985 | 11562 | 7554 | 83 |
| H81 | 9587 | 9065 | 8371 | 102 |
| H82 | 9350 | 10335 | 7858 | 102 |
| H91 | 9404 | 7133 | 7606 | 77 |
| H92 | 8760 | 6622 | 7935 | 77 |
| H101 | 8030 | 4217 | 5995 | 46 |
| H102 | 7186 | 3864 | 6126 | 46 |
| H131 | 5781 | 10456 | 8322 | 200 |
| H132 | 6184 | 12309 | 8597 | 200 |
| H133 | 6375 | 11614 | 8057 | 200 |
| H141 | 6292 | 8656 | 9166 | 197 |
| H142 | 6704 | 10479 | 9455 | 197 |
| H143 | 7209 | 8626 | 9442 | 197 |
| H151 | 6589 | 7510 | 8311 | 203 |
| H152 | 7511 | 7462 | 8573 | 203 |
| H153 | 7196 | 8620 | 8046 | 203 |
| H161 | 5302 | -1011 | 5467 | 36 |
| H201 | 4948 | 5079 | 5727 | 43 |
| H211 | 4782 | 7778 | 6246 | 51 |
| H231 | 6043 | 5199 | 7588 | 65 |
| H241 | 6228 | 2483 | 7079 | 55 |
| H251 | 5503 | 8257 | 7661 | 72 |
| H252 | 5462 | 9583 | 7164 | 72 |

| | | | | |
|---|---|---|---|---|
| H253 | 4678 | 8631 | 7264 | 72 |
| H1131 | 9090 | 6363 | 10003 | 185 |
| H1132 | 9790 | 7176 | 9763 | 185 |
| H1133 | 9069 | 6118 | 9392 | 185 |
| H1141 | 7754 | 7959 | 9811 | 184 |
| H1142 | 7577 | 9735 | 9434 | 184 |
| H1143 | 7710 | 7740 | 9196 | 184 |
| H1151 | 8935 | 9542 | 10395 | 171 |
| H1152 | 9631 | 10408 | 10161 | 171 |
| H1153 | 8777 | 11345 | 10028 | 171 |
| H2061 | 7226 | 7739 | 7512 | 68 |
| H2062 | 7145 | 9203 | 7043 | 68 |
| H2071 | 7167 | 10805 | 7843 | 99 |
| H2073 | 7924 | 11370 | 7625 | 99 |
| H2081 | 9370 | 8591 | 8475 | 100 |
| H2082 | 9240 | 10064 | 8005 | 100 |
| H2091 | 9351 | 7003 | 7670 | 85 |
| H2092 | 8621 | 6346 | 7904 | 85 |
| H2131 | 7671 | 13273 | 9835 | 198 |
| H2132 | 8190 | 11736 | 9624 | 198 |
| H2133 | 7259 | 11817 | 9397 | 198 |
| H2141 | 6984 | 15869 | 9243 | 196 |
| H2142 | 7113 | 16078 | 8659 | 196 |
| H2143 | 6561 | 14458 | 8796 | 196 |
| H2151 | 8475 | 15889 | 9556 | 203 |
| H2152 | 9009 | 14395 | 9341 | 203 |
| H2153 | 8629 | 16096 | 8978 | 203 |
| H3131 | 10268 | 15455 | 8930 | 170 |
| H3132 | 9926 | 14804 | 9421 | 170 |
| H3133 | 9338 | 15457 | 8891 | 170 |
| H3141 | 11045 | 12470 | 8906 | 186 |
| H3142 | 10715 | 11768 | 9397 | 186 |
| H3143 | 10603 | 10510 | 8882 | 186 |
| H3151 | 9611 | 14209 | 8080 | 169 |
| H3152 | 9147 | 12271 | 8013 | 169 |
| H3153 | 10083 | 12298 | 8082 | 169 |

Hydrogen coordinates (x $10^4$) and isotropic displacement parameters ($Å^2$x $10^3$) for ($R$-**2**)$_2$•D-DTTA.

**Table S6.** Torsion angles [°] for (*R*-**2**)$_2$•D-DTTA.

| | | | |
|---|---|---|---|
| C5-O1-C1-C2 | -61.9(4) | C1-O1-C5-C4 | 56.4(5) |
| C1-O1-C5-C206 | -65.4(6) | C1-O1-C5-C209 | 175.0(5) |
| C208-N201-C207-C206 | 37.6(12) | C211-N201-C207-C206 | -176.5(7) |
| C207-N201-C208-C209 | -37.0(12) | C211-N201-C208-C209 | 177.1(7) |
| C207-N201-C211-O202 | 12.6(11) | C207-N201-C211-O203 | -157.6(9) |
| C208-N201-C211-O202 | 159.9(8) | C208-N201-C211-O203 | -10.3(11) |
| O1-C1-C2-C3 | 59.3(4) | C1-C2-C3-C4 | -53.0(5) |
| C1-C2-C3-C10 | -173.7(3) | C2-C3-C4-C5 | 50.7(4) |
| C10-C3-C4-C5 | 174.2(3) | C2-C3-C10-N2 | -55.3(4) |
| C4-C3-C10-N2 | -175.8(3) | C3-C4-C5-O1 | -51.3(4) |
| C3-C4-C5-C206 | 71.6(5) | C3-C4-C5-C209 | -169.9(5) |
| C16-O4-C18-O7 | 2.6(5) | C18-O4-C16-C17 | -77.6(4) |
| C16-O4-C18-C19 | -177.2(3) | O1-C5-C206-C207 | -65.6(7) |
| C206-C5-C209-C208 | -53.6(9) | C209-C5-C206-C207 | 54.1(9) |
| O1-C5-C209-C208 | 68.0(8) | C4-C5-C206-C207 | 172.8(6) |
| C4-C5-C209-C208 | -172.9(6) | C5-C206-C207-N201 | -46.4(10) |
| N201-C208-C209-C5 | 45.3(10) | O4-C16-C17-O5 | -8.8(5) |
| O4-C16-C17-O6 | 170.1(3) | O4-C18-C19-C20 | -5.5(5) |
| O4-C18-C19-C24 | 173.6(3) | O7-C18-C19-C20 | 174.7(4) |
| O7-C18-C19-C24 | -6.2(6) | C18-C19-C20-C21 | 178.4(3) |
| C24-C19-C20-C21 | -0.7(6) | C18-C19-C24-C23 | -178.1(4) |
| C20-C19-C24-C23 | 1.0(6) | C19-C20-C21-C22 | -0.4(6) |
| C20-C21-C22-C23 | 1.0(6) | C20-C21-C22-C25 | -177.7(4) |
| C21-C22-C23-C24 | -0.6(7) | C25-C22-C23-C24 | 178.1(4) |
| C22-C23-C24-C19 | -0.4(7) | | |

Symmetry transformations used to generate equivalent atoms: #1 -x+1,y,-z+1
